# Structural insights into the evolution of ABH-fold luciferases

**DOI:** 10.1101/2025.09.13.676034

**Authors:** Marika Majerova, Jana Horackova, Jiri Damborsky, David Bednar, Martin Marek

**Author notes:** Corresponding authors: David Bednar and Martin Marek. Joint first authors (Marika Majerova, Jana Horackova).

## Abstract

The alpha/beta-hydrolase (ABH) superfamily is a widespread and functionally versatile protein fold recognized for its ability to adapt to diverse molecular functions across all three domains of life. One such spectacular example of evolutionary adaptation at the ABH-fold represents an acquisition of oxygenolytic luciferase reaction that occurred within the hydrolytic haloalkane dehalogenase family. The molecular details of this evolution remain puzzling. In this work, we determine crystal structures and explore dynamical behavior of a bifunctional ancestral ABH-fold enzyme, highlighting molecular features associated with the transition from hydrolytic to oxygenolytic catalysis at this fold. Structures showed a canonical αβα-sandwich shielded with a helical cap domain. The catalytic pocket is voluminous enough to accommodate a bulky substrate. Molecular docking showed productive binding of the coelenterazine substrate, as well as the product coelenteramide. Molecular dynamics simulations demonstrated that the coelenterazine entry does not present a major energetic barrier and identified a preferred binding orientation important for oxygenolytic catalysis. Comparisons between ancestral and extant enzymes highlighted specific amino acids and sequence motifs characteristic for oxygenolytic luciferases. Collectively, our results provide an expanded view of the evolutionary transition in which ABH-fold enzymes, originally using water to cleave chemical bonds, adapted to utilize dioxygen for bioluminescence.

## Introduction

The α/β hydrolase (ABH) fold represents a structurally adaptable scaffold that supports a broad spectrum of catalytic functions, including hydrolytic dehalogenation and oxidative bioluminescence. This fold forms the structural basis for both haloalkane dehalogenases (HLDs) (Marek et al., 2000; Newman et al., 1999; Snajdarova et al., 2023) and *Renilla*-type luciferases (Loening et al., 2007; Schenkmayerova et al., 2023), despite their drastically distinct chemistries and substrate specificities. For this reason, the ABH fold has emerged as an ideal template not only for studying the evolution of new enzymatic functions but also for protein engineering platforms aimed at expanding enzymatic function, stability, and catalytic efficiency.

The evolution of luciferase activity from a dehalogenase scaffold is thought to involve the adaptation of structural features, particularly within the cap domain and substrate-binding tunnel, to accommodate the bulkier and more hydrophobic coelenterazine (CTZ) substrate, allowing the fin ancestral sequence reconstruction (ASR) have enabled the reconstruction and engineering of proteins that bridge these two functional classes, providing snapshots of evolutionary intermediates and versatile templates for biotechnological applications (Babkova et al., 2017; Chaloupkova et al., 2019). Recent discovery of the first extant enzyme with both luciferase and HLD activity further supports the hypothesis of luminescence evolving from a dehalogenase-like ancestor (Lau et al., 2025).

Among ancestral sequences, dual-function enzymes such as Anc^HLD-RLuc^ (Chaloupkova et al., 2019) and its engineered variant Anc^INS^ (Schenkmayerova et al., 2021) have been studied for their ability to catalyze both dehalogenation and bioluminescence reactions. These proteins offer unique opportunities to dissect the molecular basis of functional divergence and to rationally engineer catalytic efficiency and stability. Another ancestral protein, DhaA 238Loc, exhibits substantially enhanced luciferase activity while retaining dehalogenase function (Musil et al., 2021). While DhaA 238Loc is similar (72.9 % identity, 84.4 % similarity) to previously reported Anc^HLD-RLuc^, its bioluminescence is improved (4.8-fold increase), but the structural determinants responsible for this improvement remain uncharacterized. Understanding how active site and aromatic core configurations contribute to the dehalogenase-to-luciferase catalysis shift is essential not only for elucidating enzyme evolution but also for advancing the rational design of improved luciferase reporters for imaging, diagnostics, and biosensing.

In this study, we determine crystal structures of DhaA 238Loc, the dual-function ancestral enzyme capable of catalyzing both hydrolytic dehalogenase and oxygenolytic luciferase activity with improved performance. Through a comprehensive structural and dynamical comparison between ancestral and extant ABH-fold enzymes, we attempt to identify and map key residues and molecular elements driving the acquisition of oxygenolytic bioluminescent reaction at this fold. Moreover, we identify residue-level adaptations that distinguish luciferase-active enzymes from the evolutionary original hydrolytic HLDs. By delineating these structural determinants, we seek to improve understanding of enzyme adaptation, inform future engineering efforts, and advance the development of enhanced bioluminescent tools.

## Materials and methods

### Protein overproduction and purification

*E*. *coli* BL21(DE3) cells were transformed with the recombinant pET21b plasmid containing DhaA 238Local gene (Musil et al., 2021) and plated on LB-agar containing 100 μg/ml ampicillin. Colonies were grown at 37 °C overnight (12–16 h). Transformed colonies were streaked into 10 mL of LB medium containing 100 μg/mL ampicillin and cultivated at 37 °C for 4 h. The entire culture was then transferred to 1 L of LB medium with 100 μg/mL ampicillin to initiate the main culture, which was grown at 37 °C for another 4 h. Enzyme expression was induced by adding IPTG (0.5 mM final concentration), followed by incubation at 20 °C for 16 h. Cells were harvested by centrifugation (4500 × g, 15 min) and resuspended in harvesting buffer (50 mM NaCl, 10 mM imidazole, 10 mM Tris, pH 7.5) before freezing at −70 °C. Prior to purification, thawed cells were treated with 90 μL DNase (1 mg/mL) and lysed by sonication (Sonic Dismembrator Model 705, Fisher Scientific, USA). The lysate was clarified by centrifugation (21036 × g, 4 °C, 1 h), and the supernatant was filtered using a 0.45 μm MilliPore syringe filter (VWR, USA). Purification of the protein was carried out in two steps. First, affinity chromatography was performed using an ÄKTA Pure system (Cytiva, USA) equipped with a 5- mL HisTrap™ excel column (Cytiva, USA). The protein was eluted with a buffer containing 50 mM NaCl, 350 mM imidazole, and 10 mM Tris (pH 7.5). In the second step, gel filtration was conducted using a HiLoad 16/600 Superdex 200 pg column (Cytiva, USA) on the ÄKTA Pure system with a buffer composed of 50 mM NaCl and 10 mM Tris (pH 7.5). Finally, the protein size and purity were verified by SDS-PAGE.

### Thermal unfolding measurements

Thermal stability and unfolding were assessed using NanoDSF Prometheus (NanoTemper, DE). The analysis was based on monitoring intrinsic tryptophan fluorescence across a temperature range of 20–95 °C with a heating rate of 1 °C/min. Measurements were conducted in triplicate, with sample concentrations of approximately 1 mg/mL. The T_m_ values were determined using ThermControl v2.0.2 software. Finally, the average values and standard deviations were calculated.

### Protein crystallization

Freshly purified protein was concentrated to approximately 10 mg/mL, and crystallization screenings were performed using a Gryphon LCP crystallization robot (Art Robbins Instruments, USA). The sitting-drop vapor-diffusion method was set up in 96-well plates (SWISSCI, CH) with 1:1 or 1:2 precipitant-to-protein ratios, and plates were incubated at 20 °C. Crystal formation was observed after 1–2 weeks. Diffraction-quality crystals were obtained in two conditions from the Morpheus screen (Molecular Dimensions, UK). First hit condition (H5 in crystallization screen) contained 0.10 M amino acids (Glutamic acid monohydrate; Alanine; Glycine; Lysine monohydrochloride; Serine - all enantiomers L and D); 0.1 M buffer (sodium HEPES and MOPS (acid)), pH = 7.5; and 30 % v/v precipitant mix (20% v/v PEG 500 MME and 10 % w/v PEG 20000). Second hit (H8 in crystallization screen) conditions were 0.10 M amino acids (Glutamic acid monohydrate; Alanine; Glycine; Lysine monohydrochloride; Serine - all enantiomers L and D); 0.1 M buffer system (sodium HEPES and MOPS (acid)), pH = 7.5, and 37.5 % v/v precipitant mix (12.5% v/v MPD, 12.5% PEG 1000 and 12.5% w/v PEG 3350). Crystals were cryo-cooled by flash freezing in liquid nitrogen for X-ray diffraction analysis.

### X-ray data collection and structure refinement

X-ray diffraction data were collected at the Swiss Light Source synchrotron using a PXIII beamline with a wavelength of 1.0 Å. Data processing was carried out using XDS (Kabsch, 2010) and scaling was performed with Aimless (Evans and Murshudov, 2013). Structures were determined by molecular replacement using Phaser (McCoy et al., 2007) with the AlphaFold2- predicted model (ColabFold v1.5.5) (Mirdita et al., 2022) as a search template. Several cycles of automatic refinement in Phenix (Liebschner et al., 2019) and manual refinement in Coot (Emsley et al., 2010) were performed. Ligands were built using eLBOW (Moriarty et al., 2009), and the final structures were graphically visualized with the PyMOL Molecular Graphics System (Version 3.1 Schrödinger, LLC).

### Bioinformatic analysis

Protein structures were aligned in the PyMOL Molecular Graphics System (Version 3.1, Schrödinger, LLC) using the following PDB IDs: 1b6g - DhlA (Ridder et al., 1999), 1mj5 - LinB (Oakley et al., 2004), 2psf - RLuc (Loening et al., 2007), 3a2m - DbjA (Prokop et al., 2010), 3u1t - DmmA (Gehret et al., 2012), 4hzg - DhaA (Lahoda et al., 2014), 4k2a - DbeA (Chaloupkova et al., 2014), 6g75 - AncHLD-RLuc (Chaloupkova et al., 2019), 6s6e - AncINS (Schenkmayerova et al., 2021), 7qxq - AncFT (Schenkmayerova et al., 2023), and 8b5o - DmmarA (Snajdarova et al., 2023); chain A was used in all cases. Protein sequences were analyzed and aligned using Clustal Omega Multiple Sequence Alignment (Madeira et al., 2024) and visualized with ESPript 3.0 (Robert and Gouet, 2014). The active site cavity volumes were calculated using FPocketWeb 1.0.1 (Kochnev and Durrant, 2022; Le Guilloux et al., 2009) and CAVER Analyst 2.0 (Jurcik et al., 2018) for each chain of the following PDB structures: 1b6g (Ridder et al., 1999), 2had – DhlA; 1mj5 (Oakley et al., 2004) , 4h7h (Okai et al., 2013) – LinB; 2psf (Loening et al., 2007), 6yn2 (Schenkmayerova et al., 2021), 7omd, 7omo, 7omr (Schenkmayerova et al., 2023) - RLuc and RLuc8; 3A2M (Prokop et al., 2010) – DbjA; 3u1t (Gehret et al., 2012) – DmmA; 4hzg (Lahoda et al., 2014), 3g9x (Stsiapanava et al., 2010) – DhaA; 4k2a (Chaloupkova et al., 2014) – DbeA; 6g75 (Chaloupkova et al., 2019) - AncHLD-RLuc; 6s6e (Schenkmayerova et al., 2021) – AncINS; 7qxq, 7qxr (Schenkmayerova et al., 2023) – AncFT; 8b5o (Snajdarova et al., 2023) - DmmarA. Calculations of root mean square deviation (RMSD) between protein structures were calculated using the Dali server (Holm et al., 2023). Figures were created in https://BioRender.com.

### Ligand preparation

The structures of CTZ (substrate) and coelenteramide (CEI, product) were prepared using Avogadro 1.2.0 software (Hanwell et al., 2012): the multiplicity of the bonds was edited to match the keto forms, all missing hydrogens were added, and the structures were minimized by the steepest descend algorithm in the Auto Optimize tool of Avogadro, using the Universal Force Field (UFF). For the molecular docking, the restrained electrostatic potential (RESP) charges of the ligands were derived by the RESP ESP charge Derive (R.E.D.) Server Development 2.0 (Vanquelef et al., 2011). Next, the AutoDock atom types were added, and PDBQT files were generated by MGLTools (Sanner, 1999; Sanner et al., 1996). For the molecular dynamics simulations, the *antechamber* module of AmberTools16 (Case et al., 2016) was used to calculate the charges of the ligands, add the atom types of the Amber force field, and compile them into PREPI parameter files. Also, the *parmchk2* tool from AmberTools16 was used to create additional FRCMOD parameter files to compensate for any missing parameters.

### Receptors preparation

The in-house solved crystal structure of DhaA 238Loc was stripped of all HETATM records (non-protein atoms), aligned to chain A of PDB ID 2psf (Loening et al., 2007) in The PyMOL Molecular Graphics System (Version 3.1 Schrödinger, LLC), and protonated with the H++ web server v. 4.0 (Anandakrishnan et al., 2012; Gordon et al., 2005), using pH = 7.5, salinity = 0.1 M, internal dielectric = 10, and external dielectric = 80 as parameters. For the subsequent docking, AutoDock atom types and Gasteiger charges were added to the receptor by MGLTools (Sanner, 1999; Sanner et al., 1996), and the corresponding PDBQT files were generated.

### Molecular docking

The AutoDock Vina 1.1.2 (Trott and Olson, 2010) software was used for molecular docking. For docking to the active site, the docking grid was specified as an x = 33.00 Å, y = 33.38 Å, z = 32.62 Å sized box, with a center in x = 47.83, y = 21.98, z = 12.41, covering the catalytic pocket and access tunnels. The flag --exhaustiveness = 100 was used to sample the possible conformational space thoroughly. The number of output conformations of the docked ligand was set to 10. The results were analyzed in The PyMOL Molecular Graphics System (Version 3.1 Schrödinger, LLC).

### System preparation and equilibration for adaptive steered molecular dynamics

DhaA 238Loc structure was prepared as described above. The crystallographic water molecules were added back into the systems, except those overlapping with the protein or ligand. Next, histidine residues were renamed according to their protonation state (HID – Nδ protonated, HIE – Nε protonated, HIP – both Nδ and Nε protonated). The ligand was prepared as described above.

Four systems were prepared for the CTZ binding analysis: four different CTZ poses were placed at the tunnel mouth of DhaA 238Loc using The PyMOL Molecular Graphics System (Version 3.1 Schrödinger, LLC). First pose was placed in the crystal-like orientation at the tunnel mouth and the others were rotated by 180° both vertically and horizontally. The *tLEaP* module of AmberTools16 (Case et al., 2016) was used to neutralize the systems with Cl^−^ and Na^+^ ions, import the ff14SB force field (Maier et al., 2015) to describe the protein and the PREPI file parameters to describe the ligand, add a truncated octahedral box of TIP3P water molecules (Jorgensen et al., 1983) to a distance of 10 Å from any atom in the system, and generate the topology, coordinate, and PDB files.

The system equilibration was carried out with the *PMEMD.CUDA* (Götz et al., 2012; Le Grand et al., 2013; Salomon-Ferrer et al., 2013) module of Amber 16 (Case et al., 2016). Five minimization steps and twelve steps of equilibration dynamics were performed. The first four minimization steps comprised 2,500 cycles of the steepest descent algorithm followed by 7,500 cycles of the conjugate gradient algorithm, while harmonic restraints gradually decreased. The restraints were applied as follows: 500 kcal·mol^−1^·Å^−2^ on all heavy atoms of the protein and ligand, and then 500, 125, and 25 kcal·mol^−1^·Å^−2^ on protein backbone atoms and ligand heavy atoms. The fifth step comprised 5,000 cycles of the steepest descent algorithm followed by 15,000 cycles of the conjugate gradient algorithm without restraint.

The equilibration MD simulations consisted of twelve steps: (i) The first step involved 20 ps of gradual heating from 0 to 298 K at constant volume using Langevin dynamics, with harmonic restraints of 200 kcal·mol^−1^·Å^−2^ on all heavy atoms of the protein and ligand, (ii) ten steps of 400 ps equilibration Langevin dynamics each at a constant temperature of 298 K and a constant pressure of 1 bar with decreasing harmonic restraints of 150, 100, 75, 50, 25, 15, 10, 5, 1, and 0.5 kcal·mol^−1^·Å^−2^ on protein backbone and ligand heavy atoms, and (iii) the last step involving 400 ps of equilibration dynamics at a constant temperature of 298 K and a constant pressure of 1 bar with no restraint. The simulations employed periodic boundary conditions based on the particle mesh Ewald method (Darden et al., 1993) for treatment of the long-range interactions beyond the 10 Å cut-off, the SHAKE algorithm (Ryckaert et al., 1977) to constrain the bonds that involve hydrogen atoms, the Berendsen barostat (Berendsen et al., 1984) at 1 bar, and the Langevin temperature equilibration using a collision frequency of 1 ps^−1^. After the equilibration, the number of Cl^−^ and Na^+^ ions needed to reach 0.1 M salinity was calculated using the average volume of the system in the last equilibration step. The whole process was repeated, from the *tLEaP* step, with the updated number of ions.

### Adaptive steered molecular dynamics

The CTZ binding trajectories were calculated using adaptive steered molecular dynamics (ASMD). The ASMD method applies to a constant external force on two atoms in the simulated systems. We simulated ligand binding by pulling two atoms together. During ASMD, several parallel simulations were started from the same state. The simulation ran in stages where the distance between the selected atoms changed by 2 Å per stage. At the end of each stage, the parallel simulations were collected and analyzed, and the Jarzynski average (Jarzynski, 1997a, 1997b) was calculated throughout the stage. The trajectory with its work value closest to the Jarzynski average was selected, and the state at the end of this trajectory was used as the starting point for the next stage. We used the default values for setting up ASMD from the respective Amber tutorial (McGee et al., n.d.) and the ASMD publication (Ozer et al., 2012).

The simulations were run with 25 parallel MDs, steered by 2 Å stages of distance increments, with a velocity of 10 Å·ns^−1^ and a force of 7.2 N. The rest of the MD settings were set as in the last equilibration step. To simulate CTZ binding, we selected the N1 atom of CTZ and the CG atom of D120 residue as steering atoms. The initial distance between the steering atoms in the equilibrated systems was measured using Measurement Wizard in The PyMOL Molecular Graphics System (Version 3.1 Schrödinger, LLC) with 2-digit precision. The selected steering atoms were pulled together to a distance of 4.4 Å, as observed in the crystal structure of azaCTZ-bound AncFT luciferase (PDB ID 7qxr) (Schenkmayerova et al., 2023).

### System preparation and equilibration for adaptive sampling

The adaptive sampling method was used to simulate product release from DhaA 238Loc with a thorough sampling of the unbinding process. The system for adaptive sampling was built using the crystal structures of DhaA 238Loc, with CEI placed in the active site, as observed in the adaptive steered MD simulation of CTZ binding.

The following steps were performed with the High Throughput Molecular Dynamics (HTMD) (Doerr et al., 2016) scripts. The protein structure was protonated with PROPKA 2.0 (Bas et al., 2008) at pH 7.5. The system was solvated in a cubic water box of TIP3P (Jorgensen et al., 1983) water molecules, with the edges at least 10 Å away from the protein by the *solvate* module of HTMD. Cl^−^ and Na^+^ ions were added to neutralize the charge of the protein and get a final salt concentration of 0.1 M. The topology of the system was built using the *amber.build* module of HTMD, with the ff14SB (Maier et al., 2015, p. 14) Amber force field and the previously compiled PREPI and FRCMOD parameter files for the ligand. The system was equilibrated using the *equilibration_v2* module of HTMD3 (Doerr et al., 2016). The system was first minimized using a conjugate-gradient method for 500 steps. Then the system was heated to 298 K and minimized as follows: (i) 500 steps (2 ps) of NVT thermalization with the Berendsen barostat with 1 kcal·mol^−1^·Å^−2^ constraints on all heavy atoms of the protein, (ii) 1,250,000 steps (5 ns) of NPT equilibration with Langevin thermostat and same constraints, and (iii) 1,250,000 steps (5 ns) of NPT equilibration with the Langevin thermostat without any constraints. During the equilibration simulations, holonomic constraints were applied to all hydrogen-heavy atom bond terms, and the mass of the hydrogen atoms was scaled by a factor of 4, enabling time steps of 4 fs (Feenstra et al., 1999; Harvey et al., 2009; Harvey and De Fabritiis, 2009; Hopkins et al., 2015). The simulation employed periodic boundary conditions, using the particle mesh Ewald method to treat interactions beyond the 9 Å cut-off. The 1-4 electrostatic interactions were scaled with a factor of 0.8333, and the smoothing and switching of van der Waals interactions were performed for a cut-off of 7.5 Å (Harvey and De Fabritiis, 2009).

### Adaptive sampling

HTMD3 (Doerr et al., 2016) was used to perform adaptive sampling of the conformations of the system. Production MD runs of 50 ns were started using the equilibrated system, employing the same settings as in the last equilibration step. The trajectories were saved every 0.1 ns. Adaptive sampling was performed using the distance of the N1 atom of CEI and the CG atom of catalytic residue D120 as the adaptive metric and time-lagged independent component analysis (tICA) (Naritomi and Fuchigami, 2013) projection in 1 dimension. 20 epochs of 10 parallel MDs were performed, corresponding to a cumulative time of 10 µs.

### Markov state model construction

The simulations were made into a simulation list using HTMD3 (Doerr et al., 2016); the water and ions were filtered out, and unsuccessful simulations shorter than 50 ns were omitted. Such filtered trajectories were combined, which resulted in 10 µs of cumulative simulation time. The CEI release was studied based on the distance of the N1 atom of CEI and the CG atom of the D120 residue. The data were clustered using the MiniBatchKmeans algorithm into 1000 clusters. Markov state models (MSM) with three macrostates were constructed at a lag time of 17 ns. The three macrostates were labelled based on the CEI location as "bound", "tunnel", and "unbound". The opening of the access tunnels was studied based on the distance of the Cα atoms of tunnel lining residue pairs: A160/L186 for the main tunnel and T155/A266 for the side tunnel. The data were clustered using the MiniBatchKmeans algorithm to 1000 clusters, and a Markov state model (MSM) with two macrostates was constructed at a lag time of 15 ns for A160/L186 (main tunnel) and 17 ns for T155/A266 (side tunnel) residue pairs.

The Chapman-Kolmogorov test was performed to ensure the models are Markovian and describe the data well (**Figures S8-9**). The states were visualized in VMD 1.9.3 (Humphrey et al., 1996), and the ligand-active site distance statistics were calculated (mean distance, SD, minimum, and maximum). The trajectory was saved for each model and visualized in The PyMOL Molecular Graphics System (Version 3.1 Schrödinger, LLC).

### Kinetic analysis using high-throughput molecular dynamics

Kinetic values (MFPT on/off, *k*_on_, *k*_off_, *k*_off_/*k*_on_, Δ*G*^0^_eq_, and *K*_D_) were calculated by the *kinetics* module of HTMD3 (Doerr et al., 2016) between the source ("unbound" state) and sink ("bound" state). Also, the equilibrium population of each macrostate was calculated and visualized. Finally, bootstrapping of the kinetics calculation was performed, using randomly selected 80 % of the data, repeated 100 times. The kinetic values were then averaged, and the standard deviations were determined.

### Binding energy calculation using MM/GBSA

The molecular mechanics/generalized Born surface area (MM/GBSA) (Genheden and Ryde, 2015; Miller et al., 2012) method was applied to calculate CEI’s binding free energy (Δ*G*_bind_) in DhaA 238Loc and the respective residue-by-residue interactions. Ante-MMPBSA.py module of AmberTools 14 (Case et al., 2016) was used to remove the solvent and ions from the original topology file, define the Born radii as mbondi3, and generate the corresponding topology files for the complex, receptor, and ligand. The Δ*G*_bind_ of CEI was calculated with the MMPBSA.py program (Miller et al., 2012). The generalized Born method was used (*&gb* namelist) with the implicit generalized Born solvent model (*igb=8*) and salt concentration 0.1 M (*saltcon=0.1*). The solvent-accessible surface area was computed with an LCPO algorithm (Weiser et al., 1999). Decomposition of the interactions (*&decomp* namelist) was generated per residue (*idecomp=2*), with discrimination of all types of energy contributions (*dec_verbose=0*). The MM/GBSA energies were calculated on 100 snapshots of each enzyme’s "bound" and "tunnel" macrostates.

### Tunnel and cavity analysis using CAVER Analyst

We used CAVER Analyst 2.0 (Jurcik et al., 2018) to calculate access tunnels in dynamic protein structures using the CAVER 3.01 algorithm (Chovancova et al., 2012; Pavelka et al., 2016). We used 1000 snapshots of the "unbound" state to analyze the tunnels leading to the active site of DhaA 238Loc and the previously studied (Horackova, 2023) RLuc8, AncFT-L14, and AncHLD-RLuc.

## Results

### Protein crystallization outcomes

We gained high-purity and stable protein, suitable for crystallization, as confirmed by the SEC elution profile and SDS-PAGE analysis (**Figure 1a,b**). NanoDSF analysis revealed a single melting transition midpoint at 70.5 ± 0.03 °C, indicating that the protein was properly folded and suitable for crystallization experiments (**Figure 1e,f**).

**Figure 1.**
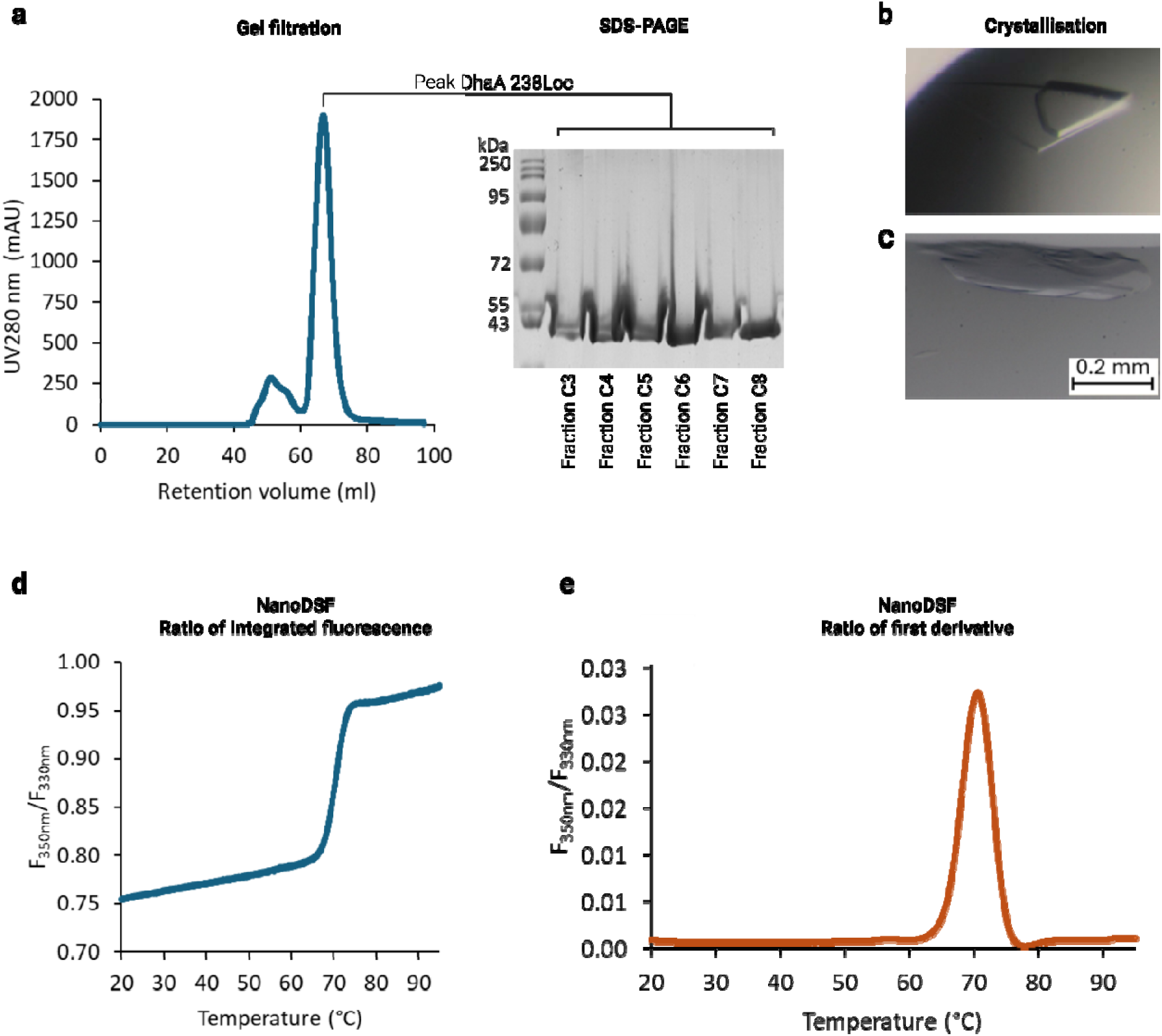
Biophysical characterization of DhaA 238Loc. **(a)** Size-exclusion chromatography elution profile, (**b**) SDS-PAGE gel of eluted peak fractions, and **(e)** integrated and **(f)** first derivative of thermal unfolding curve from NanoDSF measurements. Diffracting-quality DhaA 238Loc crystals obtained in Morpheus screen (Molecular Dimensions, UK) in well H5 (**c**) and H8 (**d**). Created in BioRender. Marek, M. (2025) https://BioRender.com/xymdl20

Many of the harvested crystals exhibited poor diffraction (>5 Å), and high-resolution data could be obtained only from crystals grown in two conditions (**Figure 1c,d**). Two complete crystallographic data sets were collected, and the protein structures were solved using molecular replacement. Iterative cycles of automated and manual refinement yielded two high-resolution structures that belonged to distinct space groups: *C*121 and *P*2_1_2_1_2 (**Table S1**). Both final models contain two enzyme molecules in the asymmetric unit. In the structure belonging to *C*121 symmetry, all residues could be built in the electron density map. Notably, additional electron density was observed in the active site and was interpreted as a polyethylene glycol (PEG) molecule. In contrast, chain B in the *P*2_1_2_1_2 structure showed region R151-W156 missing electron densities, encompassing residues in the L9 loop within the cap domain, indicating a high degree of conformational flexibility in this region (**Figure S1**). Two regions of electron densities found in chain B were interpreted es PEG molecules. Due to the higher overall model completeness, only the *C*121 structure was used for further analysis.

### Overall structure of DhaA 238Loc

Similarly to previously described HLDs (Markova et al., 2020; Schenkmayerova et al., 2021; Snajdarova et al., 2023), DhaA 238Loc features a central α/β hydrolase core domain and a typical cap domain (**Figure 2a**). The α/β hydrolase core contains an eight-stranded β-sheet, with the β2 strand oriented in an anti-parallel direction. Surrounding this central β-sheet are four α helices on one side, and three α helices on the opposite side, together forming an αβα sandwich structure. The helical cap domain is positioned on top of this sandwich, composed of four helical elements (α4, α5’, α5, α6, and α7). It is anchored between the β6 strand and the α8 helix by the L9 and L14 loops. The active site is located in a hydrophobic cavity between the α/β-hydrolase core and the cap domain.

**Figure 2.**
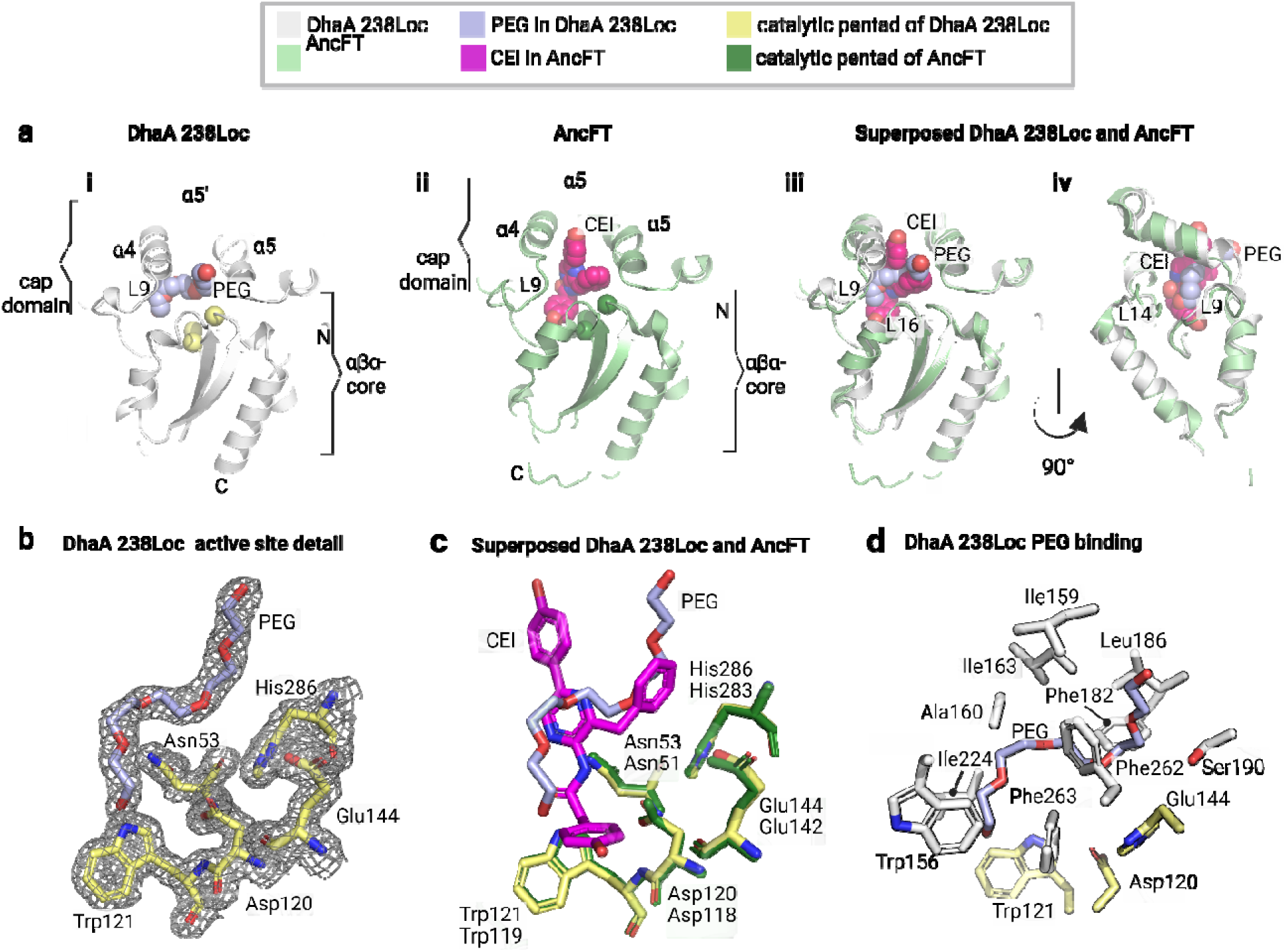
Structural comparison of DhaA 238Loc and AncFT. Crystal structure of DhaA 238Loc with bound PEG (light violet) is shown in comparison to AncFT (pale green) with bound CEI (magenta). (a) Catalytic pentad residues are highlighted as spheres: yellow in DhaA 238Loc and dark green in AncFT. Overall structures: (i) DhaA 238Loc, (ii) AncFT, (iii) front view and (iv) side view of the superposition of DhaA 238Loc and AncFT. (b) 2Fo–Fc electron density map (contour level 1σ) of the bound PEG in DhaA 238Loc, (c) superposition of the catalytic sites of DhaA 238Loc (with PEG) and AncFT (with CEI), and (d) detailed view of PEG binding in DhaA 238Loc. Created in BioRender. Marek, M. (2025) https://BioRender.com/nqon9kc

**Figure 3.**
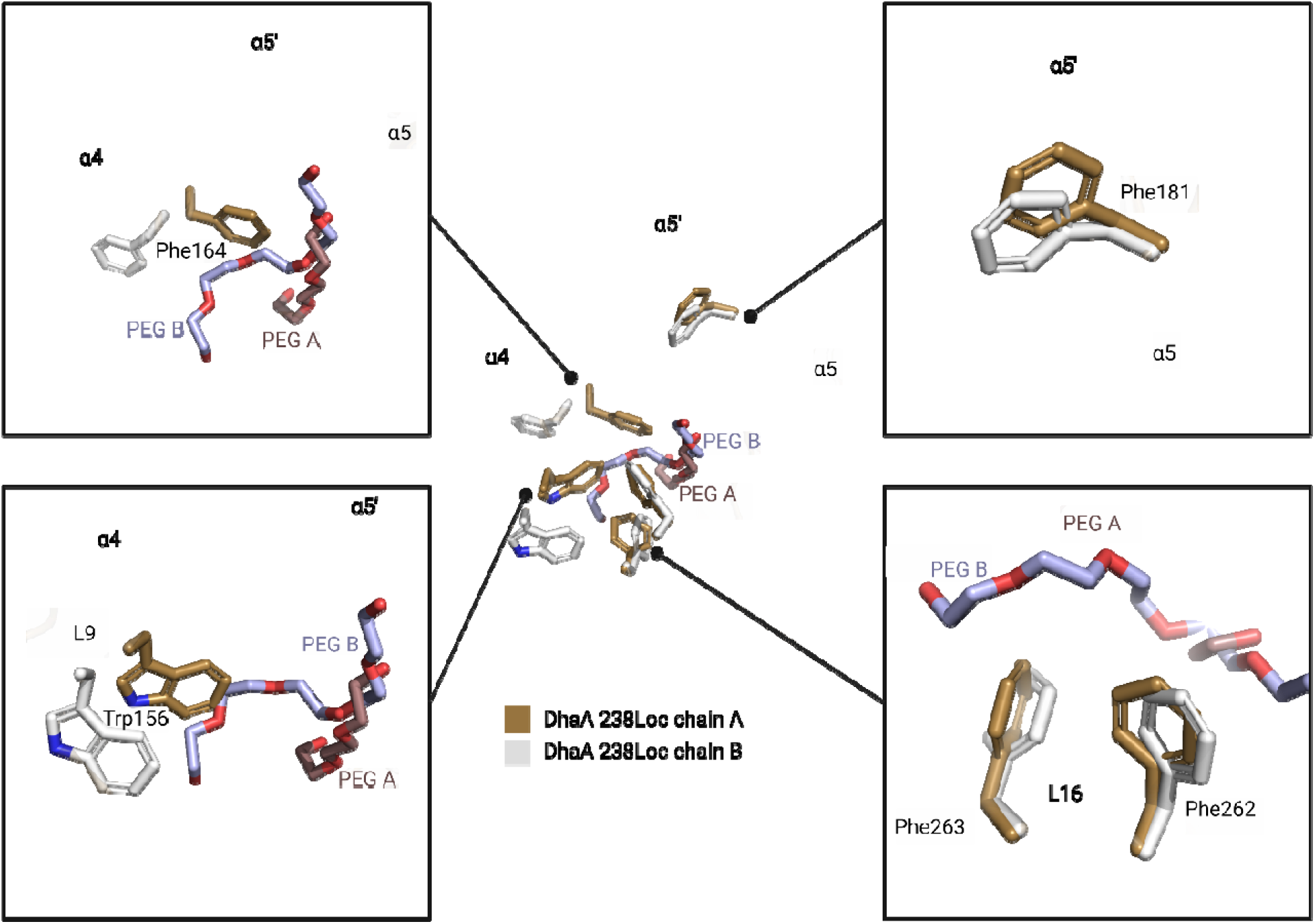
Conformational changes between DhaA 238Loc chains A and B. PEG from chain A is shown in dark violet, PEG from chain B is shown in pale violet. Created in BioRender. Marek, M. (2025) https://BioRender.com/m7a56xw

The protein structure reveals the presence of catalytic amino acids (**Figure 5b-c**). conserved within the HLD-II subfamily, including a nucleophilic aspartate (Asp120), a histidine base (His286), and a catalytic acid (Glu144), which together form a proton relay system characteristic of this enzyme family. Two halide-stabilizing residues (Asn53 and Trp121) are also present in the active site cavity. In RLuc, these residues mediate the stabilization of molecular oxygen and the leaving carbon dioxide during the catalysis. Ligand entry into the active site is likely mediated by a main tunnel located between the core and cap domains of the enzyme. In chain A, halide-stabilizing residues are involved in specific interactions with the PEG molecule, forming hydrogen bonds with its hydroxyl group at distances of 2.7 Å (Asn53) and 3.9 Å (Trp121), respectively. These residues are conserved across the enzyme family and are known to stabilize a halide ion in HLDs, or molecular oxygen and the leaving carbon dioxide in RLuc.

### Movements of aromatic residues in the catalytic pocket

The structure consists of two molecules, chains A and B (RMSD=0.8 Å), each binding a PEG molecule of different lengths and spatial arrangement. This difference is associated with a slightly different conformation, particularly involving the α4 helix, which represents the most obvious structural difference between the two chains. In this conformation, the bulky side chain of Phe164 adopts an alternative flipped-out orientation. While in chain A, Phe164 is positioned toward the active site cavity (flipped-in), in chain B, it is rotated away from the active site, creating space to accommodate the longer PEG molecule. Similarly, the phenylalanine dyad (Phe262 and Phe263), undergo rotational shifts around their side-chain axes, while Trp156, and Phe181, adopt a slightly shifted conformation. These conformational changes of bulky aromatic residues suggest a degree of plasticity in the active-site pocket that may be relevant for substrate binding, particularly for accommodating the bulky and aromatic CTZ luciferin.

### Structural adaptations toward luciferase activity

Beyond the conserved core architecture, several structural adaptations appear to distinguish oxygenolytic luciferase-active enzymes from typical HLDs. One such adaptation involves Val146, located in the L9 loop, which directly interacts with the CTZ luciferin in RLuc8. The valine dyad (V146-V147) is conserved across all compared ancestral proteins (**Figure 4b**) as well as in the known ABH-fold luciferases dafA (V149-V150) and RLuc (V146- V147) (**Figure 5**). Among the analyzed proteins, AfLuc luciferase is the only one without the valine dyad, instead, it features an M_155_C_156_ dyad at the corresponding position. (**Figure 5**). In contrast, among aligned HLDs, no sequence containing two consecutive valines is observed at this position. Specifically, DmmA and DmmAR exhibit a Leu-Val and DspA a Val-Tyr sequence, while other tested HLDs display various amino acids at these positions, such as Ile/Leu/Cys in place of Val146 and Cys/Leu/Tyr/Ala/Arg in place of Val147 (**Figure 4a**). The presence of valine residues at these positions may therefore be functionally significant, as their smaller side chain may create more space in the binding cavity, thus facilitating more effective accommodation of CTZ luciferin. In the α4 helix, Asp162 is conserved among tested luciferase sequences and some HLDs (**Figure 4a,b**). However, in the structure of DhaA 238Loc it is shifted by one position in the tertiary structure compared to RLuc8 and AncFT. At its position, isoleucine Ile163 is found instead, which interacts less effectively with the CTZ molecule. This positional shift and amino acid substitution potentially affect substrate binding and catalytic efficiency.

**Figure 4.**
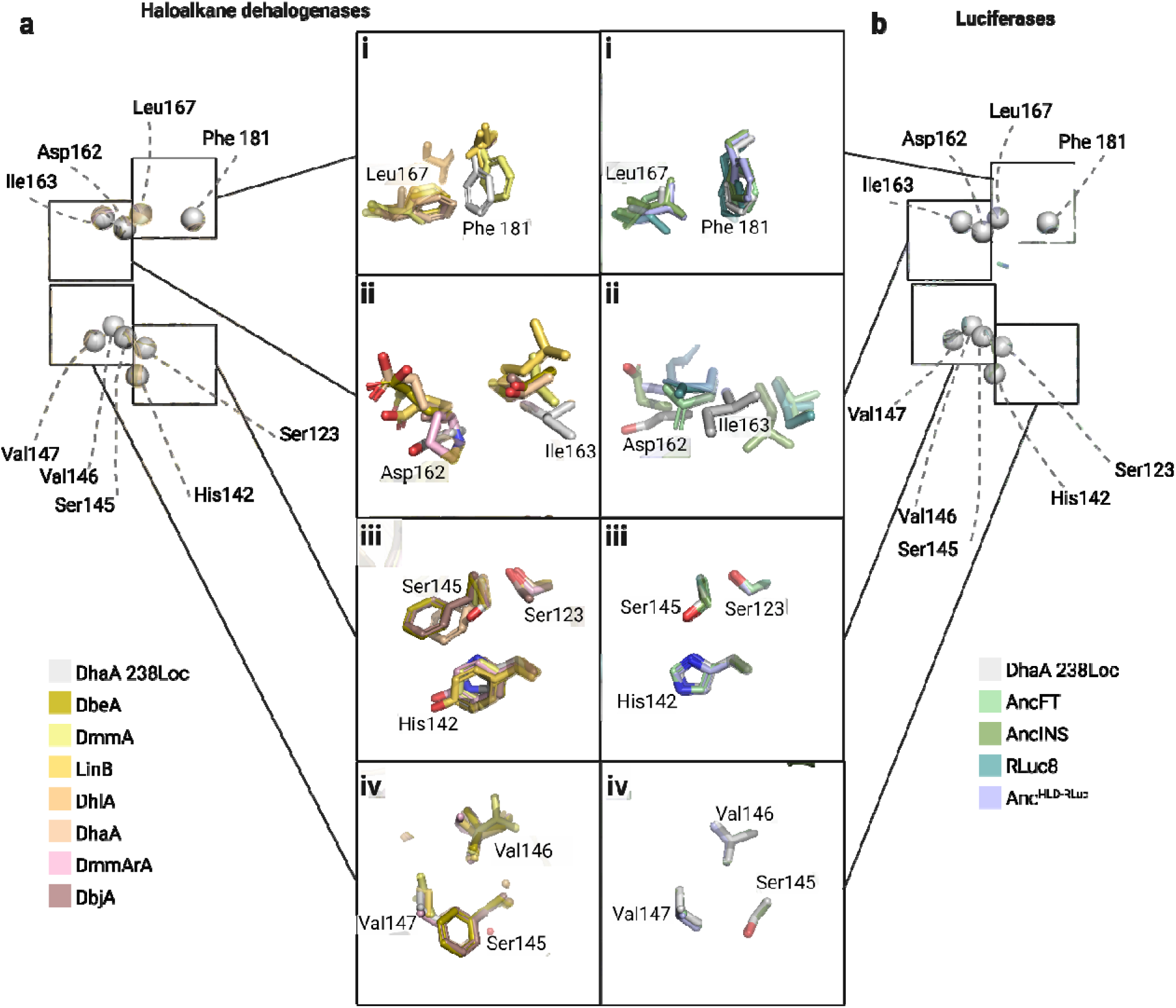
Structural comparison of key residues distinguishing oxygenolytic luciferases from hydrolytic HLDs. Residues from hydrolytic haloalkane dehalogenases are shown in shades of yellow to red (panel a), whereas residues from luciferase-active proteins are depicted in shades of blue to green (panel b). Panels (a) and (b) compare DhaA 238Loc with representative HLDs and luciferases, respectively, highlighting residues with structural adaptations linked to luciferase activity as spheres. Created in BioRender. Marek, M. (2025) https://BioRender.com/bghi6ta

Histidine at position 142 is conserved among enzymes exhibiting luciferase activity, including DhaA 238Loc, AncFT (His140), Anc^HLD-RLuc^ (His140), AncINS (His140), RLuc8 (His142), DafA (His145), and AfLuc (His151). In contrast, typical HLDs, except for DspA, possess hydrophobic residues at this position, such as Ile, Tyr, Phe, or Cys. The presence of polar histidine may contribute to local structural rearrangements, potentially facilitating the opening of the active site necessary for accommodating the CTZ substrate.

Similarly, Ser145 is conserved among enzymes displaying luciferase activity. In its surroundings, another serine can be found, Ser123. These serine residues, both hydrophilic in nature, may contribute to increased local flexibility and potentially facilitate active site opening, particularly when acting together with His142. In contrast, in HLDs, the Ser145 position is typically occupied by apolar or hydrophobic residues. This composition forms a compact hydrophobic core in the α/β hydrolase sandwich. DspA contains glycine at this position (Gly145), while DhlA, LinB, DmmA, and DmmArA possess alanine (position 149, 133m 169 and 120, respectively), and DhaA, DbjA, and DbeA have phenylalanine (position 131, 128 and 128, respectively). The environment surrounding this region in HLDs is predominantly hydrophobic, consisting of neighboring hydrophobic or aromatic residues (residue numbers refer to DhaA238Loc in alignment) : Phe/Tyr/Cys at position 142, Phe127 (or Leu131 in DhlA and Met151 in DmmA), Ile/Val/Gly/Leu/Phe at position 270, Ile/Leu/Val at position 140, Phe/Tyr/Arg/Leu at position 255, and Trp/Cys/Val/Leu at position 274.

Leu167 is a conserved residue among proteins exhibiting luciferase activity, where either leucine or isoleucine is consistently found at this position (**Figure 4b and Figure 5**). Similarly, DspA, DmmA, and DmmaR also contain leucine at this position (Leu166, Leu194 and Leu140, respectively). Phenylalanine occupies the equivalent site in LinB (Phe154), DhaA (Phe152), DbeA (Phe151), and DbjA (Phe160), while in DhlA, it is replaced by tryptophan (Trp175). In its proximity, Phe181 is conserved at the equivalent position in all compared proteins with luciferase activity, except for AfLuc, where it is substituted by tyrosine (Tyr190). Among HLDs, with the exception of DmmA, the Phe181 is generally replaced with smaller non-aromatic residues. In other HLDs, this position is typically occupied by Glu, Met, Val, or Ala.

**Figure 5.**
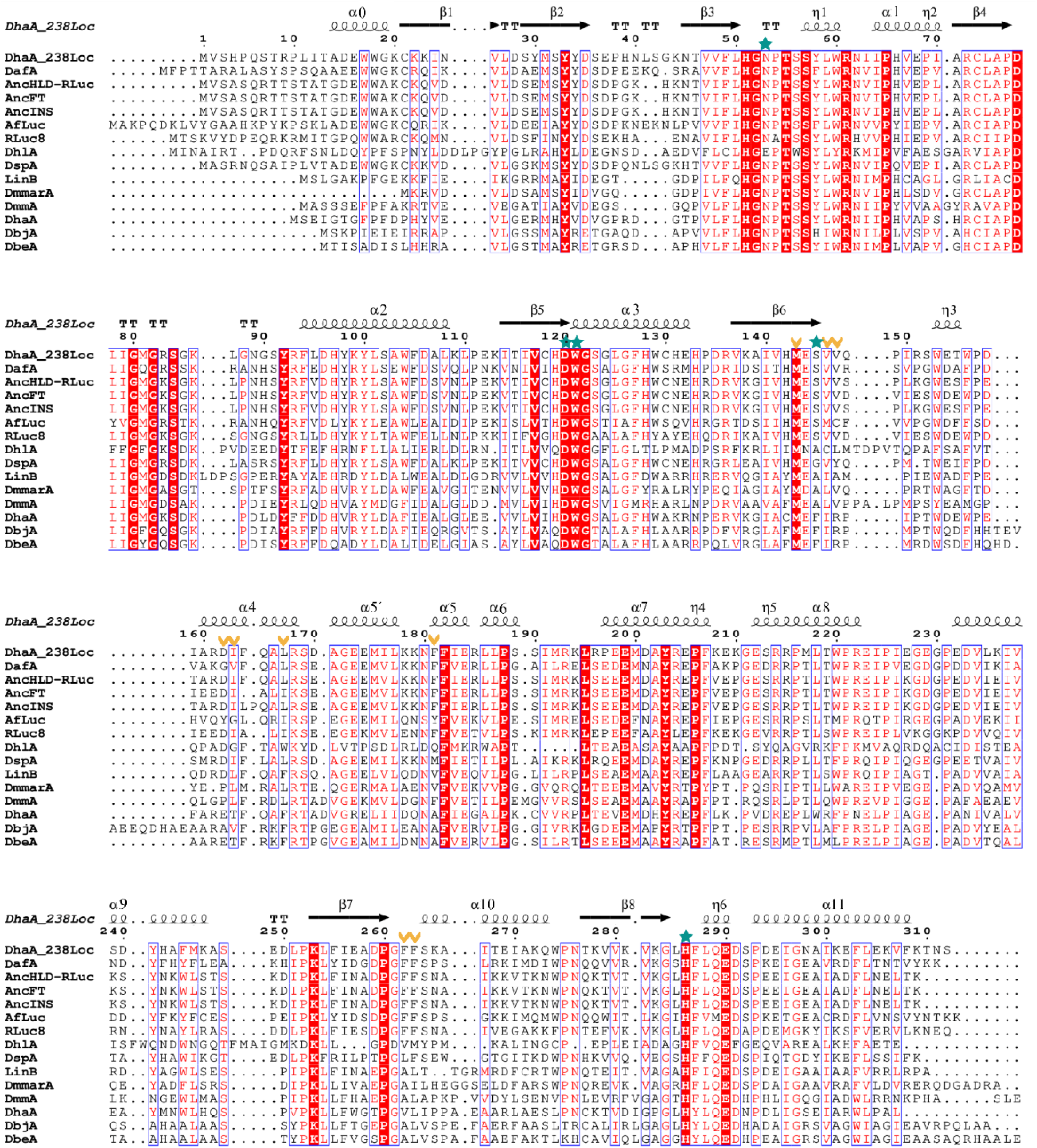
Multiple sequence protein alignment of DhaA 238Loc and selected proteins: HLDs (LinB, DhlA, DspA, DhaA, DbjA, DbeA, DmmarA), luciferases (RLuc8, AfLuc) and dual function enzyme (DafA, Anc^HLD-RLuc^, AncFT, AncIns). Amino acids of the putative conserved active site are marked with green stars, amino acids different between enzymes with only luciferase activity and HLD/dual function enzymes are marked with yellow arrows. Created in BioRender. Marek, M. (2025) https://BioRender.com/bjgdr9u

### Presence of the aromatic core

The tightly packed cluster of aromatic residues lines the substrate-binding pocket and plays a role in shaping the microenvironment necessary for efficient catalysis and substrate stabilization. Position equivalent to 156 in DhaA238Loc varies between luciferases and HLDs and may influence the substrate access path (**Figure 6a,b**). In DhaA 238Loc, AncFT, RLuc, and DhaA, this position is occupied by a tryptophan. In contrast, AncHLD-RLuc, AncINS, dafA, AfLuc, DhlA, DspA, LinB, DmmAR, DbjA, and DbeA all feature a phenylalanine at the same site. Interestingly, DmmA differs from both groups, lacking tryptophan or phenylalanine at this position and instead containing a methionine (Met183), reflecting a divergent sequence in this region. These substitutions likely affect the physicochemical properties and shape of the entrance tunnel, potentially influencing substrate binding dynamics.

**Figure 6.**
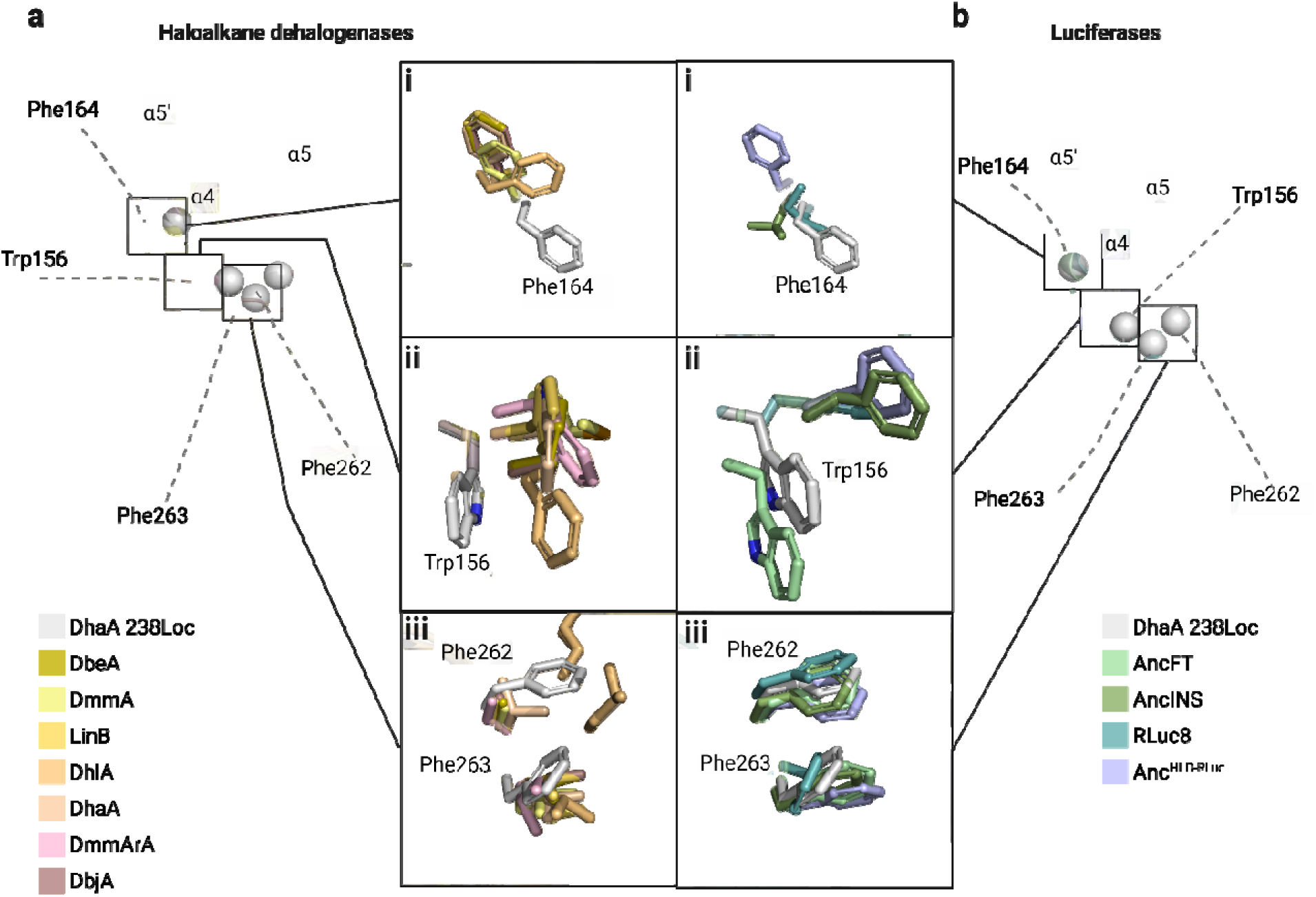
Structural comparison of aromatic core residues differing between luciferases and HLDs. Residues from hydrolytic haloalkane dehalogenases are shown in shades of yellow to red (panel a), whereas residues from luciferase-active proteins are depicted in shades of blue to green (panel b). Panels (a) and (b) compare DhaA 238Loc with representative HLDs and luciferases, respectively, highlighting residues of the aromatic core as spheres. Created in BioRender. Marek, M. (2025) https://BioRender.com/abue6b3

Another bulky residue, Phe164, is absent in the active sites of efficient luciferases such as AncFT, AncINS, RLuc8, and AfLuc. Instead, these proteins feature smaller residues at the corresponding position: Ile163 in AncFT, Leu162 in AncINS, Ile161 in RLuc (**Figure 6b**), and Leu173 in AfLuc. In contrast, phenylalanine at equivalent positions is present in HLDs and in ancestral sequences with low luciferase activity, suggesting a potential link between the presence of a bulky phenylalanine residue and reduced catalytic efficiency for luciferase activity. An exception is DmmarA, which does not contain phenylalanine at this position but instead carries a methionine (Met137), a residue with a long side chain, occupying the same structural location. Similarly, DhlA retains phenylalanine at this position (Phe172), adopting a conformation comparable to that observed in DhaA 238Loc (**Figure 6a**).

Crucially, the phenylalanine dyad, Phe262 and Phe263 in DhaA 238Loc, is conserved among all oxygenolytic proteins exhibiting luciferase activity (**Figure 5**), indicating their functional importance in stabilizing the active site architecture and facilitating substrate binding. In contrast, these positions vary considerably among HLDs. For instance, DhlA features Val262- Met263, DspA has Leu_261_Phe, and DmmarA contains Ala_232_Ile. DhaA harbours Val245-Leu246, while DbeA, DbjA, DmmA, and LinB all possess the Ala–Leu combination. The unique and consistent presence of two bulky aromatic residues in luciferases, compared to the smaller or less aromatic residues in HLDs, likely contributes to forming a more compact and hydrophobic microenvironment essential for efficient CTZ binding and bioluminescent activity. Interestingly, in chain B of the DhaA 238Loc structure, Phe263 adopts a distinct conformation compared to the other analyzed structures. Its side chain is rotated inward, facing into the active site cavity. Thi conformational change correlates with the presence of a PEG molecule bound in chain B, in contrast to the shorter PEG observed in chain A. A similar orientation of Phe263 is seen only in the RLuc8 structure (PDB ID 7omo, Schenkmayerova et al., 2023).

### Voluminous enzymatic pocket accommodates bulky luciferin

All structural features and differences described above collectively contribute to differences in the active site cavity volumes. When comparing these volumes, the following trend is observed where oxygenolytic CTZ-utilizing ABH-fold luciferases generally have larger active site cavities than HLDs (**Figure 7g-i**). This molecular adaptation reflects the need to accommodate the bulkier luciferin for the bioluminescent reaction. The increased cavity size may support more efficient substrate binding and the conformational dynamics required for the light- emitting reaction and luciferin/oxyluciferin (un)binding events. Notable exceptions to this trend are DbjA and DmmA, two HLDs that display cavity volumes comparable to luciferases (Figure 7g-i), suggesting either latent structural potential or inherent flexibility despite their lack of luciferase activity.

**Figure 7.**
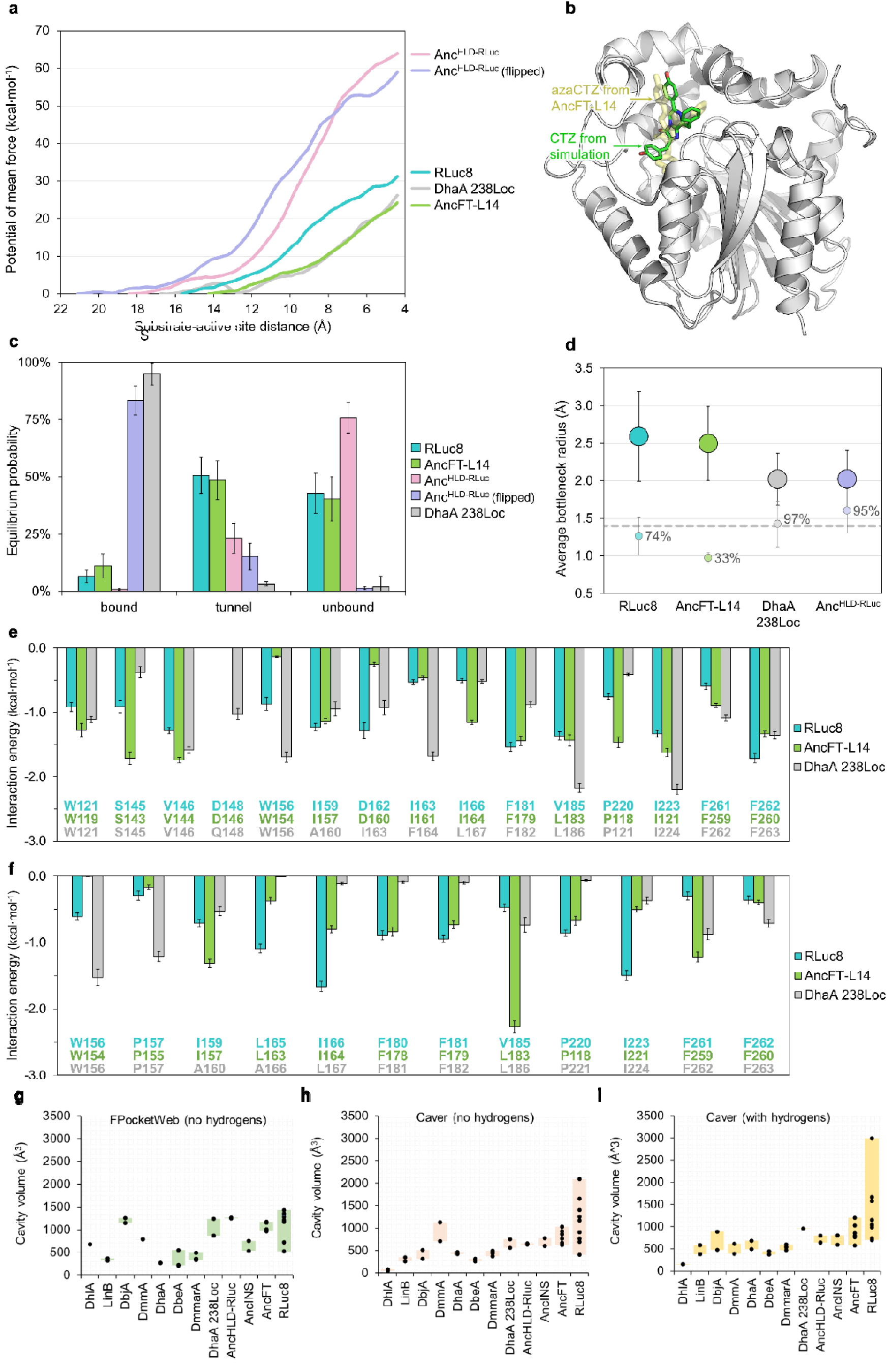
MD-based analysis and comparison of DhaA 238Loc with RLuc8, AncFT-L14, and Anc^HLD-RLuc^. **(a)** The potential of mean force used to pull CTZ into the active site of luciferases obtained by adaptive steered MD. **(b)** CTZ end-pose (green sticks) from the simulation of binding to DhaA 238Loc (white cartoon), showing a crystallographic ligand as reference (translucent pale-yellow sticks, PDB ID 7ome). **(c)** The equilibrium probabilities of coelenteramide (CEI) macrostates based on the CEI-active site distance ("bound", "tunnel", "unbound") from the adaptive sampling of CEI unbinding. The simulations were started from the crystal-like CEI pose for each enzyme, plus from the flipped pose in Anc^HLD-RLuc^. The error bars indicate the standard deviation of the equilibrium probability from bootstrapping a random 80 % of the data 100 times. **(d)** Analysis of access tunnels with CAVER Analyst on 1000 dynamic snapshots of luciferase structures. Main tunnels, shown as the bigger circles, have wider average bottlenecks and were present in over 99 % of snapshots. In comparison, the side tunnels are shown as smaller, lighter circles, have narrower average bottlenecks, and are less common (percentages next to the circles). **(e, f)** The interaction energies of CEI per residue of (e) the "bound" CEI macrostate and (f) the "tunnel" macrostate from adaptive sampling with RLuc8, AncFT-L14, and DhaA 238Loc. For clarity, only attractive interactions are shown (<−1.0 kcal·mol^-1^ for the "bound" states and <−0.5 kcal·mol^-1^ for the "tunnel" states). The error bars indicate the standard error. **(g, h, i)** Comparison of active-site cavity volumes in DhaA 238Loc, luciferase RLuc8, dual-function ancestral proteins (Anc^HLD-RLuc^, AncINS, AncFT), and HLDs (DhlA, LinB, DbjA, DmmA, DbeA, DmmarA), calculated using FPocketWeb **(g)**, CAVER Analyst with hydrogen atoms **(h)**, and without hydrogen atoms **(i)**. Created in BioRender. Marek, M. (2025) https://BioRender.com/c6xzq3l

### Plausible binding poses revealed by molecular docking

To further probe the functional relevance of the observed cavity architecture, we performed molecular docking of the luciferase substrate CTZ and its product CEI to DhaA 238Loc. The docked poses were compared with known structures of luciferase-active enzymes such as RLuc8, AncFT-L14, and Anc^HLD-RLuc^ (Horackova, 2023). While the predicted poses in RLuc8 and AncFT-L14 closely resembled the crystallographic luciferins in PDB ID 7ome (azaCTZ) and 7qxq (CEI, Schenkmayerova et al., 2023), the poses in DhaA 238Loc slightly differ. However, they still fit in the catalytic site and have reasonable energies, unlike those in Anc^HLD-RLuc^ (**Figures S3-4, Table S3**). Interestingly, DhaA 238Loc has ∼1 kcal·mol^−1^ higher affinity towards the product than to the substrate, which is the opposite for RLuc8 and AncFT- L14 and could point towards slow product unbinding or product inhibition in DhaA 238Loc. Notably, we were docking to fixed protein structures, which influenced the docking results. We used the “open” state crystal structures with a more spacious active site cavity. To supplement the docking results, we used CAVER Analyst 2.0 (Jurcik et al., 2018) to calculate the cavity volumes for “open” and “closed” states on the available crystal structures The active site cavity in DhaA 238Loc is more spacious than AncHLD-RLuc but still smaller than in RLuc8 (**Figure S5**).

### Substrate binding is energetically favorable

We simulated substrate binding using adaptive steered MD (ASMD). Among four orientations tested (**Figure S6**), pose 1, resembling the CTZ orientation in the structure of luciferase (PDB ID 7ome) was the most accessible (**Figure 7b)**, consistent with prior results for RLuc8 and AncFT-L14. Notably, the substrate binds most easily to AncFT-L14, closely followed by DhaA 238Loc (1.9 kcal·mol^−1^ difference), while RLuc required 5 kcal·mol^−1^ more work than DhaA 238Loc (**Figure 7a**). In contrast, CTZ binding to Anc^HLD-RLuc^ required almost twice as much work as to RLuc8, and the preferred starting pose was “flipped” compared to the crystal pose (**Figure S7**). These results suggest that limited activity in DhaA238Loc is unlikely due to geometrically unfavorable substrate binding or high energy barrier.

### Product release is hampered

CEI release was studied using adaptive sampling MD from both the crystallographic pose and the “flipped” Anc^HLD-RLuc^ pose. Notably, the crystal-like pose of CEI in Anc^HLD-RLuc^ was not observed experimentally nor predicted by docking; therefore, it was created artificially by transplanting CEI from the crystal structure of AncFT (PDB ID 7qxq). The simulations were clustered to separate the "bound" CEI state, the "tunnel" state (CEI in the access tunnel), and the "unbound" CEI state (**Figures S8-9, Table S3**), and to calculate their equilibrium probabilities (**Figure 7c**), and estimate kinetics (**Table S4**).

As reported previously by Toul and co-workers (Toul et al., 2025), he simulations have indicated that the product release is energetically favorable in RLuc8 and AncFT-L14, as evidenced by a "bound" state with low equilibrium probability and positive ΔG, *k_off_*faster than *k_on_*; **Figure 7c, Tables S3-4**. On the other hand, the result of the simulations with Anc^HLD-RLuc^ depended heavily on the product’s orientation: the crystal-like pose followed the same trend as RLuc8 and AncFT- L14 with a stronger preference for the “unbound” state, while the energetically preferred flipped pose followed the opposite trend, explaining previously observed product inhibition (Schenkmayerova et al., 2021). Interestingly, in DhaA 238Loc, we also observed strong product binding in the active site (“bound” state having a 95% equilibrium probability and ΔG of -2.56 kcal·mol ¹, indicating slower product release.

In the "bound" macrostates, the product mainly interacted with the same residues in RLuc8 and AncFT-L14, except for Trp156 and Asp162, which lacked a significant affinity in AncFT-L14 (**Figure 8e**). The interactions in DhaA 238Loc were also quite similar to RLuc8, the main differences being a lower affinity of the product to Ser145, a higher affinity to Trp156, Phe163 (corresponding to Ile163 in RLuc8), and Pro220, and a significant interaction with Gln148 (Asp148 in RLuc8), which was not detected in RLuc8 or AncFT-L14.

However, the "tunnel" macrostates showed key differences (**Figure S9**). In RLuc8, the product interacted with the dynamical L9/α4 fragment (**Figure S10**) during release, while in a more rigid AncFT-L14, it left straight through the main tunnel. In contrast, in the simulation with DhaA 238Loc, the product leaves through the side (slot) tunnel, which is opened due to the dynamic loop L9. Therefore, the per-residue interactions differ noticeably (**Figure 7f**). In RLuc8, the product interacts mainly with Leu165, Ile166, Phe180, Phe181, Pro220, and Ile223; in AncFT-L14 with Ile159, Ile166, Phe180, Phe181, Leu185 (Val185 in RLuc8), and Phe261; while in DhaA 238Loc with Trp156, Pro157, Phe261, and Phe262, which are the residues surrounding the side tunnel.

## Anatomies of enzyme access tunnels

Analysis of access tunnels using CAVER Analyst 2.0 across 1000 “unbound” snapshots showed that all four studied enzymes maintained a main tunnel and a side tunnel (**Figure 7d**). RLuc8 and AncFT-L14 had wider main tunnels (average bottleneck radii 2.6 Å and 2.5 Å), while DhaA238Loc and AncHLD-RLuc were narrower (both 2.0 Å). Side tunnels were more frequently present in DhaA238Loc and AncHLD-RLuc (97% and 95%), and less in RLuc8 and AncFT-L14 (75% and 33%), with corresponding bottleneck radii of 1.43, 1.60, 1.27, and 0.98 Å, respectively. These results fit well with the “tunnel” state in which the ligand interacts mainly with residues forming the slot tunnel rather than the main tunnel (**Figure S9** and **Figure 7f**).

## Discussion

In this study, we present structural and dynamical insights into the bifunctional ancestral enzyme DhaA 238Loc, which exhibits dual dehalogenase-luciferase catalysis. DhaA 238Loc adopts the canonical ABH fold, characteristic of both HLDs and *Renilla*-type luciferases. It features a central β-sheet flanked by α-helices, and a cap domain that forms a large active site cavity. PEG molecule, originating from the crystallization mother liquor, bound within the protein cavity, illustrates the accessibility and ligand-accommodating nature of this pocket.

The presence of dual activity in DhaA 238Loc supports the hypothesis that *Renilla*-type luciferases evolved from ABH-fold ancestors with dehalogenase activity (Delroisse et al., 2021, 2017; Lau et al., 2025). Similarly, a recent study demonstrated the evolutionary adaptation of a distinct hydrolytic ABH-fold enzyme, 1-H-3-hydroxy-4-oxoquinaldine 2,4-dioxygenase, to perform O -dependent oxygenolytic catalysis. The work highlighted how evolution repurposed multiple structural elements of the ABH fold to enable the transition from hydrolytic to oxygenolytic function (Bui et al., 2023). Comparison of DhaA 238Loc to extant *Renilla* luciferase (RLuc8), ancestral proteins with luciferase activity (AncHLD-RLuc, AncINS, AncFT), and HLDs (LinB, DhaA, DbjA, DmmA, DmmarA, DhlA) reveal several shared and divergent structural features. One of the key determinants of luciferase activity and spectral properties in ABH-fold enzymes is the position and role of residue Asp162 at the rim of the active site cavity. In RLuc8 and AncFT, this aspartate (Asp160) plays a critical role in modulating the electronic state of the CEI product. Specifically, it acts as a nucleophile, stabilizing the 6-p- hydroxyphenolate anion of CEI and fine-tuning the emission wavelength (Schenkmayerova et al., 2023). However, the equivalent Asp162 residue in DhaA 238Loc is rotated outward, and its role appears compromised. Its position is instead occupied by Ile163, potentially reducing the stabilization of a negatively charged CEI emitter. This structural rearrangement suggests a partial loss of the spectral-tuning mechanism present in the more specialized luciferases.

In addition to Asp162, DhaA 238Loc features an aromatic cluster formed by Phe262 and Phe263, the same π-stacking residues that line the CTZ-binding pocket in RLuc8. These residues, rare in typical dehalogenases, likely help stabilize the substrate via hydrophobic and π– π interactions, facilitating luciferase activity (Schenkmayerova et al., 2023). The stabilizing effect of Phe262 and Phe263 on luciferin was confirmed by computing interaction energies from MD simulation data. A comparable π–π stacking pattern is observed in AncFT, where the 8- benzyl group of azaCTZ interacts with Phe259 and Phe260, while the hydroxyl group of the 6- (p-hydroxyphenyl) moiety forms a hydrogen bond with Asp160. Such interactions highlight the critical synergy between aromatic residues and catalytic side chains in stabilizing high-energy intermediates and fine-tuning color of emitted light.

Furthermore, residues in the L9 and L16 loops, notably Trp153, Trp156, Phe261, and Phe262 in RLuc8, have been shown to regulate active site openness and impact activity, with mutagenesis studies confirming a 2- to 3-fold decrease in luminescence (Schenkmayerova et al., 2023). DhaA 238Loc partially mirrors this architecture, especially with bulky aromatic residues in corresponding loops, suggesting increased flexibility and substrate accessibility compared to canonical dehalogenases. Additionally, the Ser143 backbone carbonyl (equivalent to RLuc8) is positioned to form a hydrogen bond with CEI, potentially contributing to the transient stabilization of the emitter state.

To place DhaA 238Loc in a functional context, we used molecular modelling to compare its substrate binding and product release to RLuc8, AncFT-L14, and Anc^HLD□RLuc^. ASMD simulations showed that luciferin binding to DhaA 238Loc is not more difficult than in the two specialized luciferases, RLuc8 and AncFT-L14, unlike Anc^HLD□RLuc^, which has the greatest energetic barrier. Additionally, molecular docking revealed that DhaA 238Loc slightly prefers its product over the substrate, opposite to RLuc8 and AncFT-L14, which suggests a tendency toward slower unbinding or product inhibition. Adaptive sampling MD of product release confirmed that the product remains tightly bound in DhaA 238Loc, whereas RLuc8 and AncFT-L14 release CEI more readily. Together, these findings indicate that substrate access is not a limiting factor in DhaA 238Loc, but slow product release likely lowers its overall turnover relative to modern luciferases.

Together, these structural and dynamical features point toward an evolutionary transition in DhaA 238Loc, where luciferase activity is enabled through partial specialization of the binding pocket, even without a full optimization for light emission.

## Acknowledgments

The authors would like to express their thanks to the Czech Ministry of Education, Youth and Sports (TEAMING CZ - CZ.02.01.01/00/23_029/0008437-, EXCELES Neuro - LX22NPO5107, ESFRI RECETOX - LM2023069) and Czech Science Foundation (GA22-09853S and GX25-17329X). This project has received funding from the European Union’s Horizon 2020 research and innovation programme under grant agreement No. 857560 TEAMING, and by the European Union Centre of Excellence CLARA (No. 101136607). The article reflects the author’s view, and the Agency is not responsible for any use that may be made of the information it contains. We acknowledge CF Biomolecular Interactions and Crystallography of CIISB, Instruct-CZ Centre, supported by MEYS CR (LM2023042) and European Regional Development Fund-Project „Innovation of Czech Infrastructure for Integrative Structural Biology“ (No. CZ.02.01.01/00/23_015/0008175). Computational resources were provided by the e-INFRA CZ and ELIXIR-CZ projects (90254 and LM2023055), supported by the Ministry of Education, Youth and Sports of the Czech Republic. The authors are thankful to the Swiss Light Source (SLS) synchrotron members for using their beamline facilities and their help during data collection. The text was revised for clarity and grammar using ChatGPT (OpenAI). All scientific content and interpretations were conceived and verified by the authors.

## Author contributions

M.Maj. and J.H. contributed equally to this work. M.Maj. prepared protein samples, carried out the crystallization screenings, and optimized crystallization hits. M.Maj. and M.M. collected diffraction data and solved the protein crystal structure. J.H. and D.B. performed molecular docking and MD and ASMD simulations, respectively. J.D., D.B., and M.M. designed the project, supervised research, and interpreted data. M.Maj., J.H. and M.M. wrote the manuscript with the contribution of all authors. All authors have approved the final version of the manuscript.

