## Supplementary file for "Structural insights into the evolution of ABH-fold luciferases"

[*]Joint first authors

**Table S1**. Data collection and refinement statistics.

| Data collection* | DhaA 238Loc (*C*121) | DhaA 238Loc ($P2_12_12$) |
|---|---|---|
| Wavelength (Å) | 1 | 1 |
| Resolution range | 46.43 – 1.598 (1.656 – 1.598) | 46.84 – 2.199 (2.277 – 2.199) |
| Space group | *C*121 | $P2_12_12$ |
| Unit cell parameters a, b, c (Å) | 83.501 81.773 91.726 | 78.643 81.325 93.689 |
| Unit cell parameters α, β, γ (°) | 90 99.574 90 | 90 90 90 |
| Total reflections | 540,674 | 413,426 |
| Unique reflections | 79,229 | 31,185 |
| Multiplicity | 6.8 | 13.3 |
| Completeness (%) | 98.57 (96.5) | 99.69 (98.69) |
| Mean I/sigma (I) | 17.03 (1.72) | 12.13 (0.99) |
| Wilson B-factor | 20.94 | 41.1 |
| R-merge | 0.062 (1.009) | 0.198 (2.87) |
| CC1/2 | 0.999 (0.746) | 0.999 (0.446) |
| Reflections used in refinement | 79,229 | 31,184 |
| Reflections used for R-free | 3,977 | 1,493 |
| R-work | 0.159 (0.292) | 0.232 (0.377) |
| R-free | 0.198 (0.334) | 0.29 (0.431) |
| Number of non-hydrogen atoms | 5,638 | 4,968 |
| Macromolecules | 5,032 | 4,830 |
| Ligands | 46 | 20 |
| Solvent | 560 | 118 |
| Protein residues | 607 | 592 |
| RMS (bonds) | 0.014 | 0.007 |
| RMS (angles) | 1.56 | 1.16 |
| Ramachandran favored (%) | 96.19 | 91.13 |
| Ramachandran allowed (%) | 3.81 | 7.68 |
| Ramachandran outliers (%) | 0 | 1.19 |
| Rotamer outliers (%) | 0.55 | 0.19 |
| Clash score | 3.08 | 8.42 |
| Average B-factor | 29.29 | 58.39 |
| macromolecules | 28.01 | 58.6 |
| ligands | 53.71 | 64.3 |
| solvent | 38.82 | 49.04 |
| PDB ID | 9RBN | 9RBP |

*Statistics for the highest-resolution shell are shown in parentheses.

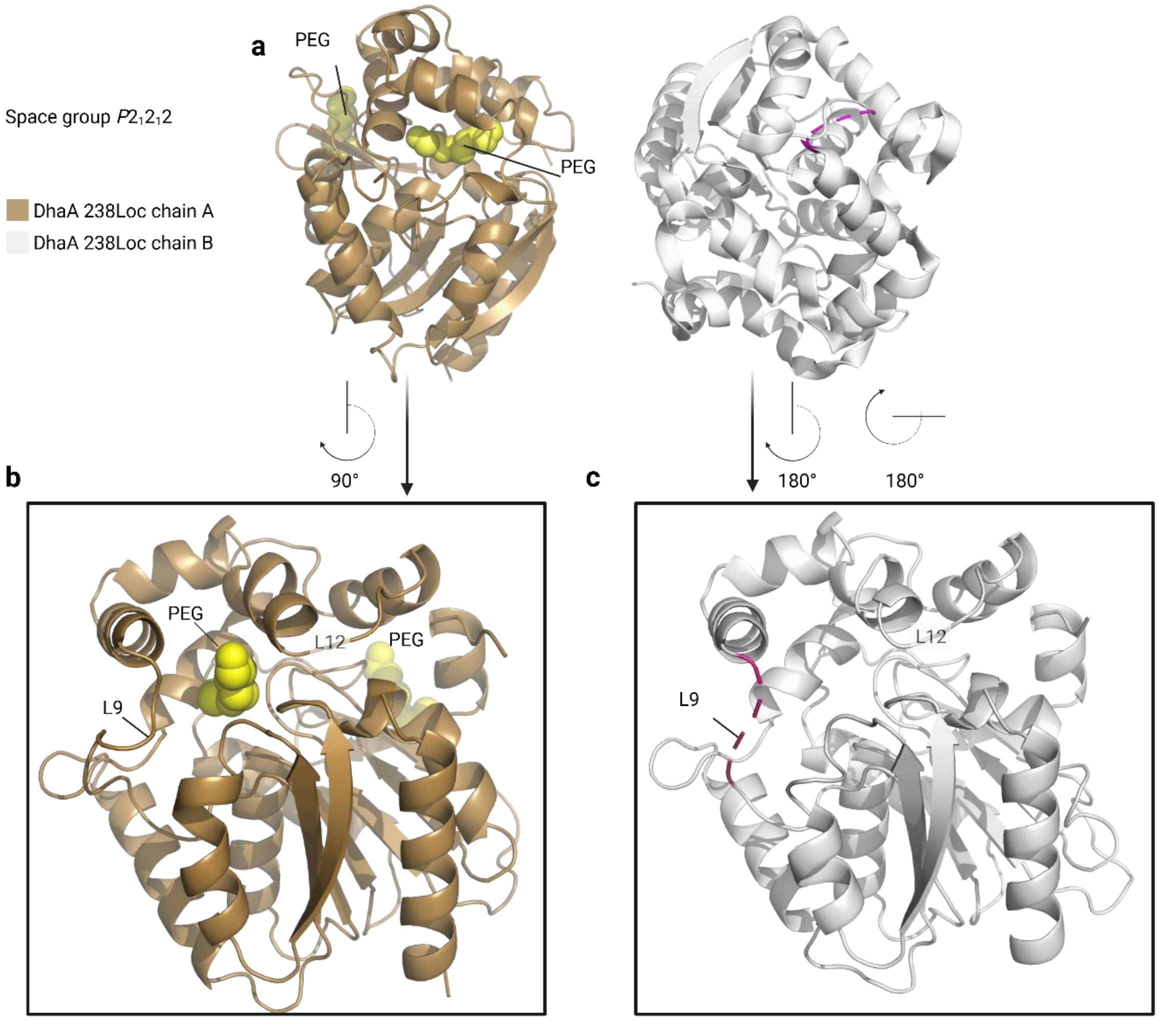


**Figure S1**: Crystal structure of DhaA 238Loc, space group *P2₁2₁2*. (**a**) Assymetric unit containing chain A (sand) and B (grey) (**b**) detail of chain A and (**c**) chain B with missing electron density (magenta) in Loop 10. Created in BioRender. Marek, M. (2025) https://BioRender.com/6naywwf

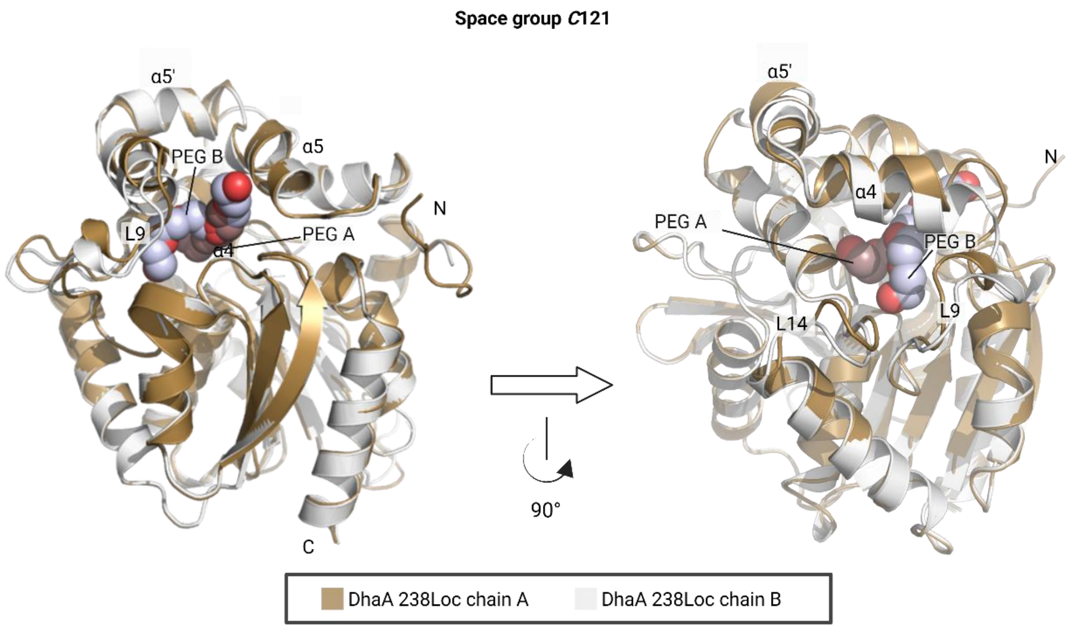


**Figure S2:** Comparison of DhaA 238Loc chain A and chain B, space group *C121.* (a) front view and (b) side view. Created in BioRender. Marek, M. (2025) https://BioRender.com/v7pqdag

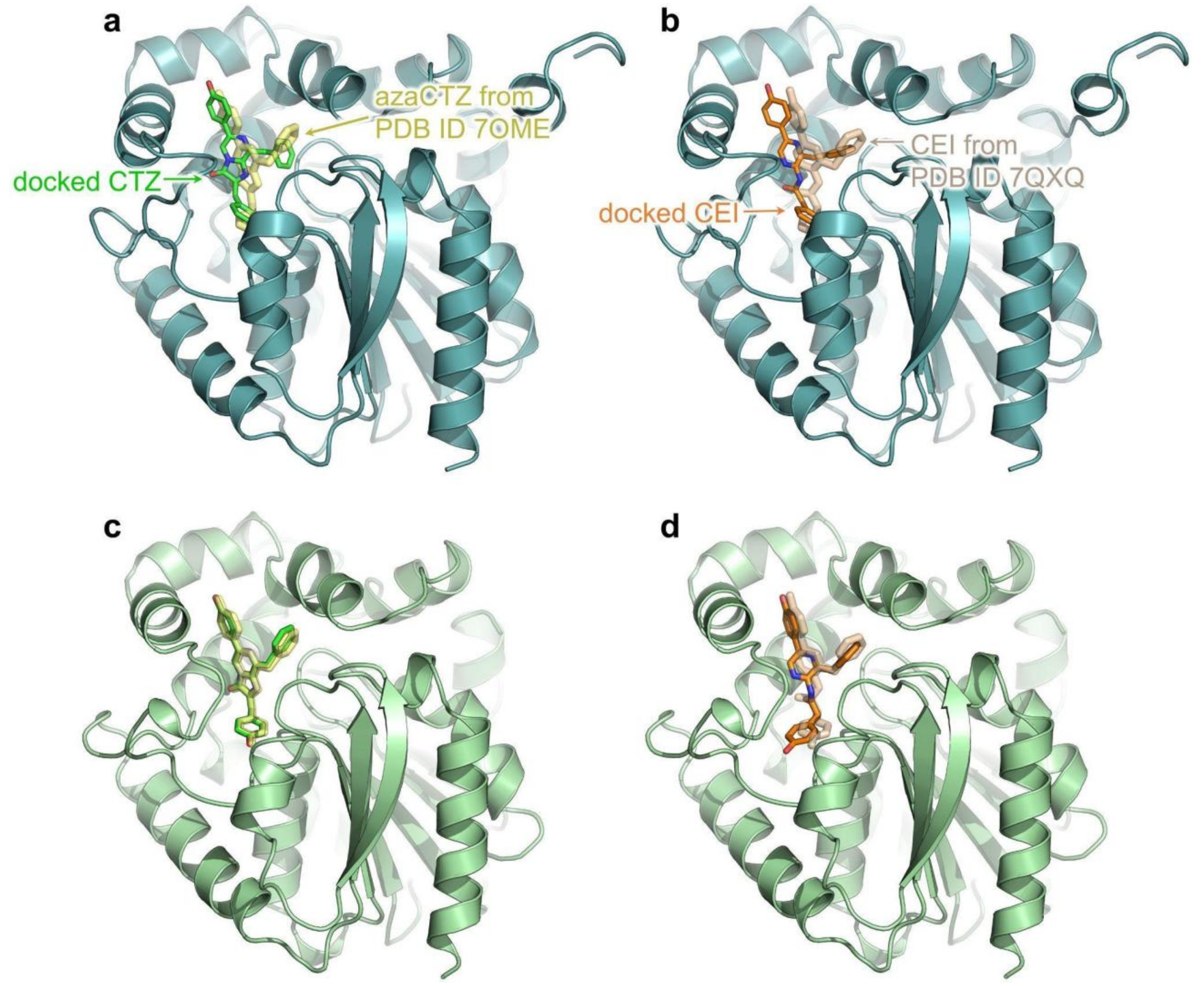


**Figure S3:** Molecular docking of coelenterazine (CTZ) and coelenteramide (CEI) to the structure of RLuc8 (PDB ID 2psf) shown as a light teal cartoon (a,b) and to the structure of AncFT-L14 (PDB ID 7ome) shown as a pale green cartoon (c,d). The best predicted poses of CTZ are shown as green sticks in (a,c), while the crystallographic pose from 7ome is represented by transparent pale yellow sticks. The best predicted poses of CEI are shown as orange sticks in (b,d), while the crystallographic pose from 7qxq is shown as transparent wheat-colored sticks. Structure 7ome depicts the azaCTZ-bound AncFT-L14 luciferase, and 7qxq the CEI-bound AncFT luciferase.

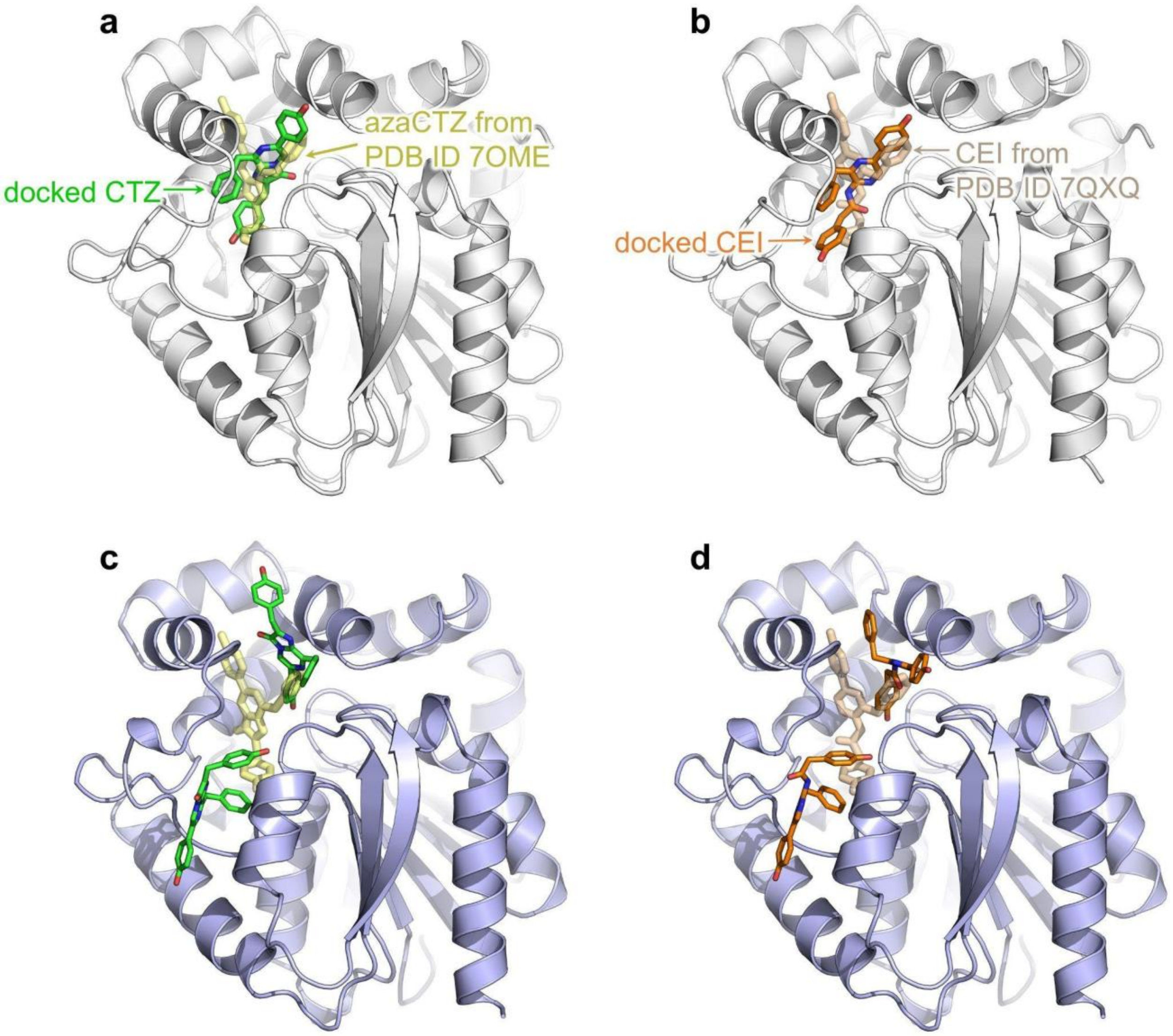


**Figure S4:** Molecular docking of coelenterazine (CTZ) and coelenteramide (CEI) to the structure of DhaA 238Loc (in-house structure) shown as a white cartoon (a,b) and to the structure of $Anc^{HLD-RLuc}$ (PDB ID 6g75) shown as a light blue cartoon (c,d). The best predicted poses of CTZ are shown as green sticks in (a,c), while the crystallographic pose from 7ome is represented by transparent pale yellow sticks. The best predicted poses of CEI are shown as orange sticks in (b,d), while the crystallographic pose from 7qxq is shown as transparent wheat-colored sticks. Structure 7ome depicts the azaCTZ-bound AncFT-L14 luciferase, and 7qxq the CEI-bound AncFT luciferase.

**Table S2:** Predicted affinities of coelenterazine (CTZ) and coelenteramide (CEI) from molecular docking to RLuc8, AncFT-L14, DhaA 238Loc, and $Anc^{HLD-RLuc}$.

| protein | ligand affinity (kcal·mol$^{-1}$) |
| --- | --- |

| | CTZ | CEI |
|---|---|---|
| **RLuc8** | -12.1 | -11.3 |
| **AncFT-L14** | -13.2 | -12.2 |
| **DhaA 238Loc** | -9.6 | -10.8 |
| **Anc$^{\textbf{HLD-RLuc}}$** | -9.3 | -9.3 |

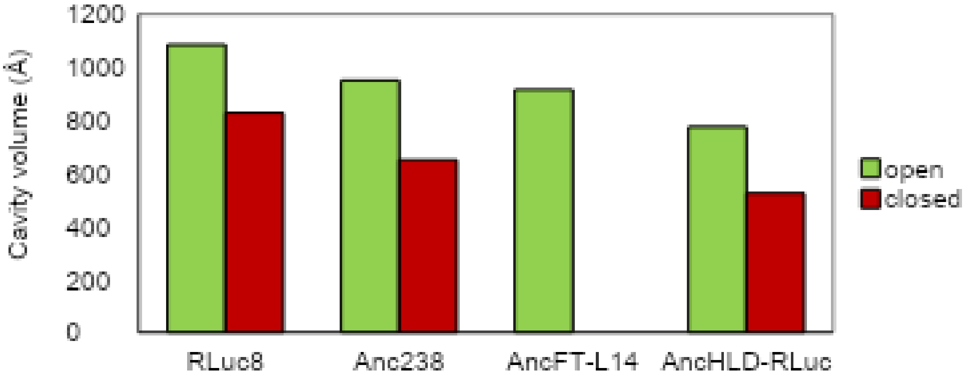


**Figure S5:** The active site cavity volumes in the open and closed crystal structures of RLuc8 (PDB ID 2psf), DhaA 238Loc (in-house structure), AncFT-L14 (PDB ID 7ome), and Anc$^{HLD\text{-}RLuc}$ (PDB ID 6g75) computed in CAVER Analyst 2.0. For AncFT-L14, only one chain is available (open structure).

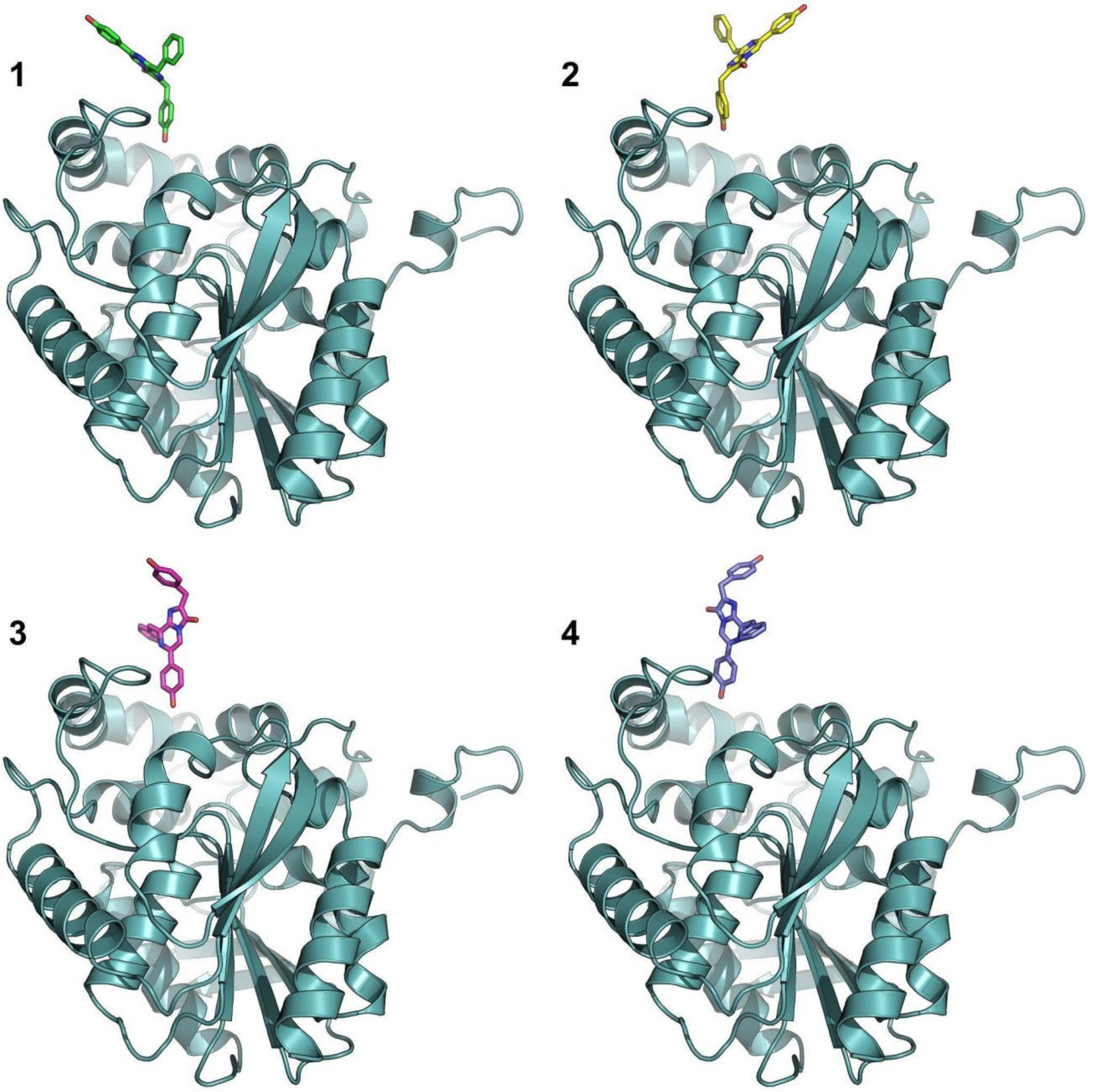


**Figure S6:** Starting poses of CTZ for the simulation of substrate binding. CTZ poses 1-4 are shown as sticks in green, yellow, magenta, and slate colours, respectively. The RLuc8 structure is shown as a light teal cartoon.

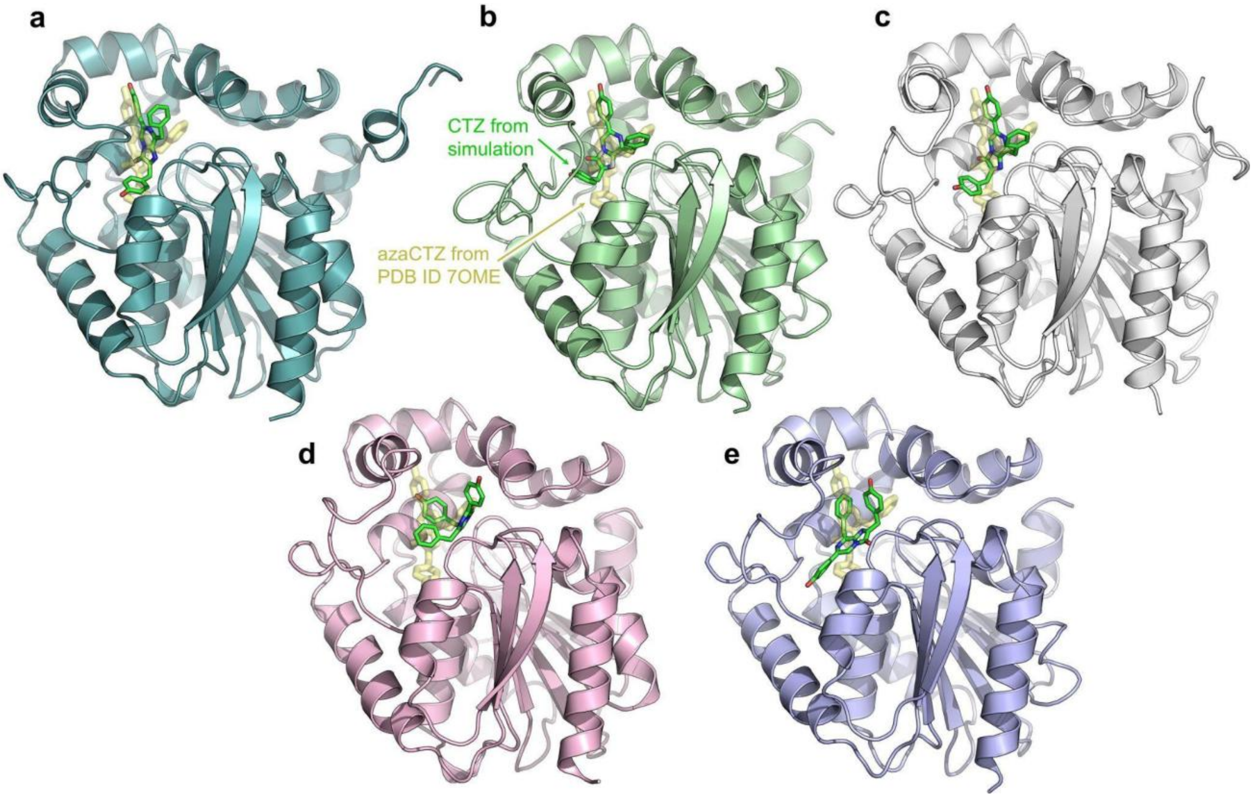


**Figure S7:** Bound coelenterazine (CTZ, green sticks) from binding simulations started from pose 1 (**Figure S4**) using the ASMD method. The binding was simulated in (a) the RLuc8 structure (light teal cartoon), (b) AncFT-L14 (pale green), (c) DhaA 238Loc (white), and (d) $Anc^{HLD\text{-}RLuc}$. Additionally, image (e) shows the result of CTZ binding to $Anc^{HLD\text{-}RLuc}$ from pose 3. For reference, the crystallographic azaCTZ from the AncFT-L14 crystal structure (PDB ID 7ome) is shown as transparent pale-yellow sticks.

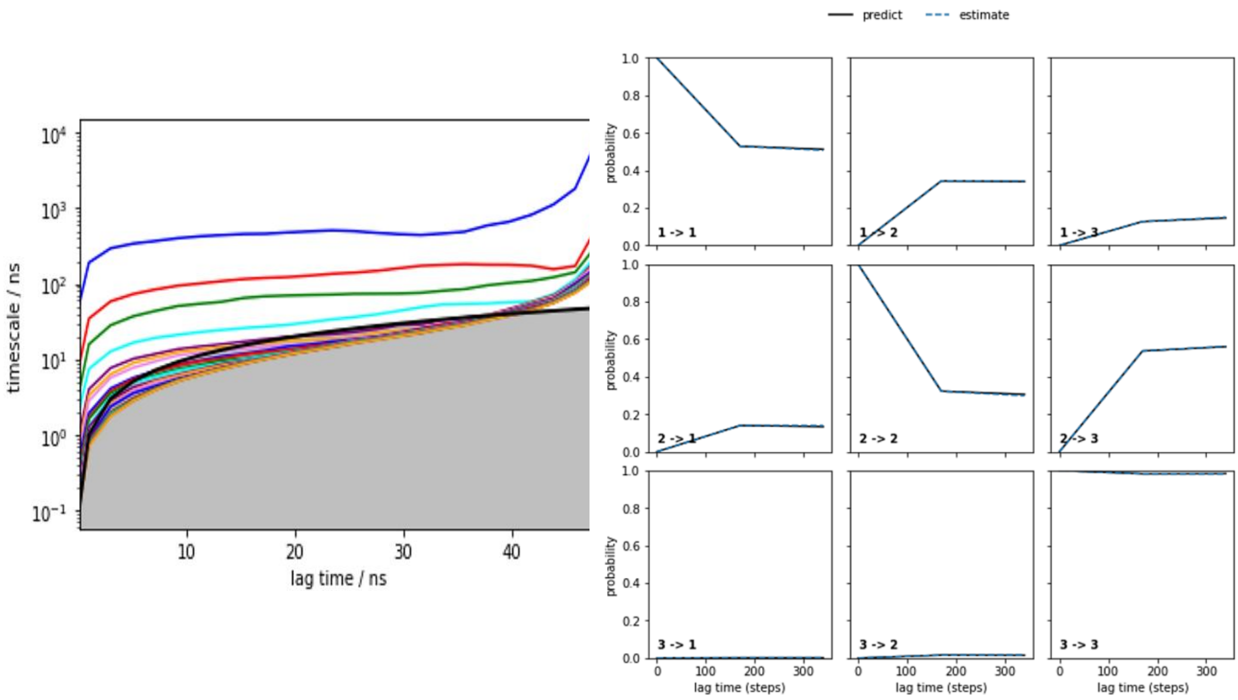


**Figure S8:** Implied timescales and Chapman-Kolmogorov test of MSMs of CEI release from DhaA 238Loc. The implied timescales (left) show the transitions between the macrostates. The Chapman-Kolmogorov test (right) was done for a Markov state model of three macrostates constructed at a 17ns lag time. The selected lag time is the lowest at which the “predict” and “estimate” lines agree.

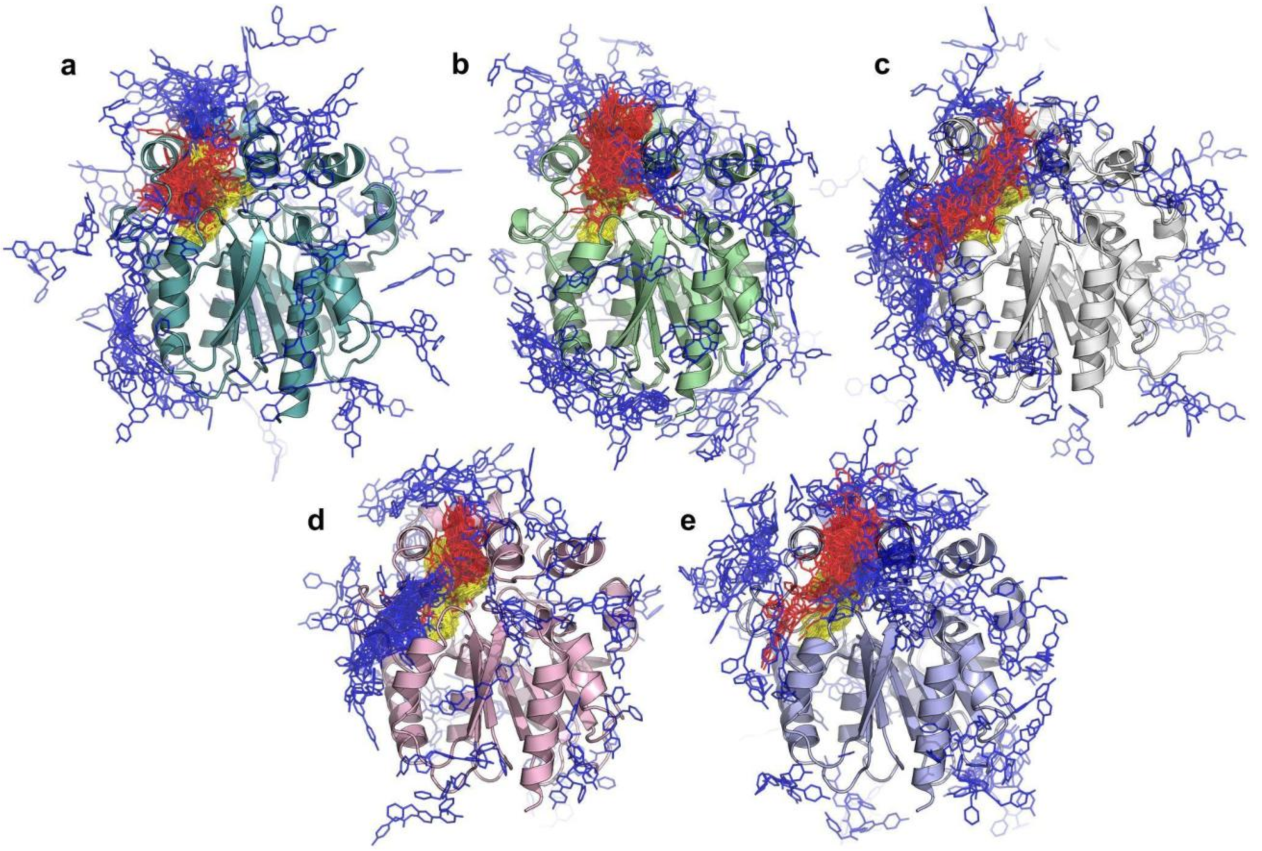


**Figure S9:** Coelenteramide (CEI) macrostates from Markov state models (MSMs) from adaptive sampling of CEI unbinding from (a) RLuc8 (light teal cartoon), (b) AncFT-L14 (pale green), (c) DhaA 238Loc (white), (e) $Anc^{HLD\text{-}RLuc}$ (light pink), and (f) flipped CEI unbinding from $Anc^{HLD\text{-}RLuc}$ (light blue). The macrostates are shown as thin sticks and colored based on their distance to the active site (yellow, red, and blue in ascending order of distance).

**Table S3:** Summary of the mean CEI-active site distance and the equilibrium probabilities of macrostates from Markov state models from the adaptive sampling of CEI release from RLuc8, AncFT-L14, DhaA 238Loc, and $\text{Anc}^{\text{HLD-RLuc}}$.

| system | macrostate | mean distance (Å) | eq. probability |
|---|---|---|---|
| | "bound" | 5.17 ± 0.16* | 0.066 ± 0.028 |
| RLuc8 | "tunnel" | 10.79 ± 0.34 | 0.506 ± 0.079 |
| | "unbound" | 28.16 ± 0.31 | 0.428 ± 0.088 |
| | "bound" | 4.59 ± 0.38 | 0.111 ± 0.052 |
| AncFT-L14 | "tunnel" | 11.22 ± 0.87 | 0.485 ± 0.085 |
| | "unbound" | 28.22 ± 0.50 | 0.404 ± 0.096 |
| | "bound" | 6.56 ± 0.19 | 0.949 ± 0.048 |
| DhaA 238Loc | "tunnel" | 14.42 ± 0.86 | 0.033 ± 0.010 |
| | "unbound" | 28.36 ± 1.04 | 0.019 ± 0.047 |
| | "bound" | 4.81 ± 0.07 | 0.009 ± 0.004 |
| $\text{Anc}^{\text{HLD-RLuc}}$ | "tunnel" | 9.99 ± 0.11 | 0.232 ± 0.066 |
| | "unbound" | 25.00 ± 0.30 | 0.759 ± 0.067 |
| | "bound" | 6.11 ± 0.07 | 0.834 ± 0.063 |
| $\text{Anc}^{\text{HLD-RLuc}}$ (flipped CEI) | "tunnel" | 14.43 ± 0.23 | 0.153 ± 0.058 |
| | "unbound" | 29.66 ± 0.46 | 0.014 ± 0.006 |

*The values were obtained from bootstrapping a random 80 % of the data 100 times.

**Table S4:** Coelenteramide (CEI) unbinding kinetics from adaptive sampling simulations with RLuc8, AncFT-L14, DhaA 238Loc, and $\text{Anc}^{\text{HLD-RLuc}}$. The kinetic parameters were calculated between the "bound" and "unbound" Markov state models.

| **enzyme** | **RLuc8** | **AncFT-L14** | **DhaA 238Loc** | **$\text{Anc}^{\text{HLD-RLuc}}$** | |
|---|---|---|---|---|---|
| **pose** | **crystal-like CEI** | | | | **flipped CEI** |
| **$\Delta G^*$ (kcal·mol$^{-1}$)** | 1.15 ± 0.31** | 0.81 ± 0.36 | -2.56 ± 0.41 | 2.72 ± 0.34 | -2.33 ± 0.40 |
| **$k_{on}$ (M$^{-1}$·s$^{-1}$)** | (2.68±0.80)×10$^5$ | (2.17±0.73)×10$^5$ | (2.15±0.30)×10$^6$ | (2.13±0.84)×10$^4$ | (4.50±0.91)×10$^5$ |
| **$k_{off}$ (s$^{-1}$)** | (1.14±0.22)×10$^6$ | (7.09±1.74)×10$^5$ | (3.59±1.65)×10$^4$ | (1.26±0.19)×10$^6$ | (3.86±1.38)×10$^4$ |
| **$k_{off}/k_{on}$ (M)** | 5.12 ± 3.92 | 3.88 ± 2.14 | 0.02 ± 0.01 | 84.35 ± 143.53 | 0.09 ± 0.05 |
| **$K_D$ (M)** | 8.15 ± 5.91 | 4.61 ± 2.63 | 0.02 ± 0.10 | 137.37 ± 305.44 | 0.02 ± 0.01 |

*ΔG – the energy of the "bound" state (negative ΔG = stable “bound” state); $k_{on}$ – rate constant of binding; $k_{off}$ – rate constant of unbinding; $k_{off}/k_{on}$ – the ratio between $k_{off}$ and $k_{on}$; $K_D$ – dissociation constant of the "bound" state

**The values were obtained from bootstrapping a random 80 % of the data 100 times.

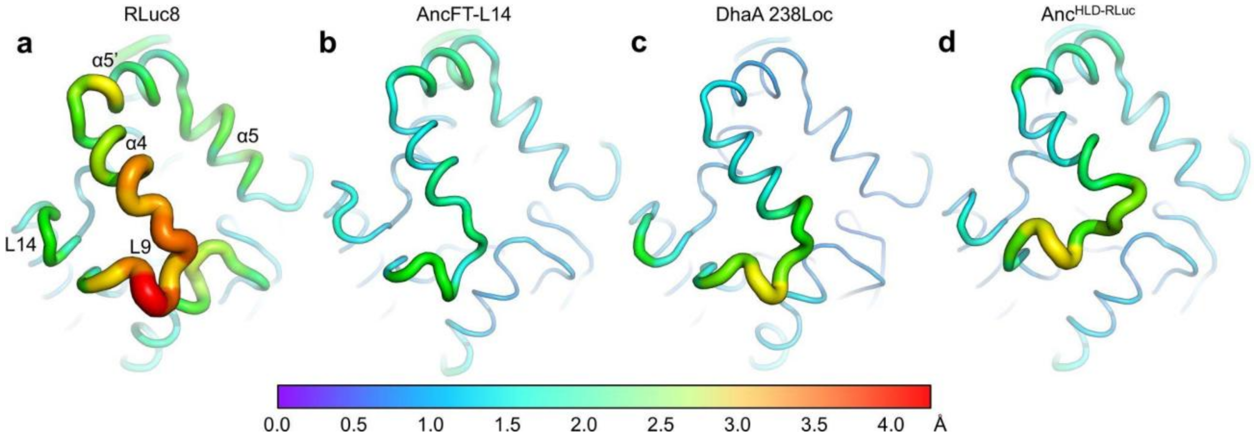


**Figure S10:** The dynamics of the cap domains of (a) RLuc8, (b) AncFT-L14, (c) DhaA 238Loc, and (d) $Anc^{HLD-RLuc}$. The root-mean-square fluctuations (RMSF) of Cα atoms in the "unbound" state from adaptive sampling were mapped onto the structures, shown as the thickness and color of the backbone structure representation.
